# The conformational landscape of fold-switcher KaiB is tuned to the circadian rhythm timescale

**DOI:** 10.1101/2024.06.03.597139

**Authors:** Hannah K. Wayment-Steele, Renee Otten, Warintra Pitsawong, Adedolapo M. Ojoawo, Andrew Glaser, Logan A. Calderone, Dorothee Kern

## Abstract

How can a single protein domain encode a conformational landscape with multiple stably-folded states, and how do those states interconvert? Here, we use real-time and relaxation-dispersion NMR to characterize the conformational landscape of the circadian rhythm protein KaiB from *Rhodobacter sphaeroides*. Unique among known natural metamorphic proteins, this KaiB variant spontaneously interconverts between two monomeric states: the “Ground” and “Fold-switched” (FS) state. KaiB in its FS state interacts with multiple binding partners, including the central KaiC protein, to regulate circadian rhythms. We find that KaiB itself takes hours to interconvert between the Ground and FS state, underscoring the ability of a single sequence to encode the slow process needed for function. We reveal the rate-limiting step between the Ground and FS state is the *cis-trans* isomerization of three prolines in the fold-switching region by demonstrating interconversion acceleration by the prolyl isomerase CypA. The interconversion proceeds through a “partially disordered” (PD) state, where the C-terminal half becomes disordered while the N-terminal half remains stably folded. We discovered two additional properties of KaiB’s landscape. Firstly, the Ground state experiences cold denaturation: at 4°C, the PD state becomes the majorly populated state. Secondly, the Ground state exchanges with a fourth state, the “Enigma” state, on the millisecond timescale. We combine AlphaFold2-based predictions and NMR chemical shift predictions to predict this “Enigma” state is a beta-strand register shift that eases buried charged residues, and support this structure experimentally. These results provide mechanistic insight in how evolution can design a single sequence that achieves specific timing needed for its function.

**Significance Statement:** One can conceptualize KaiB as an on-off switch to regulate circadian rhythms in bacteria, where the “On state” is the Fold-switched state that binds KaiC and other proteins, and the “Off state” is the Ground state. Our work exemplifies how evolution tuned the kinetics of interconversion to align with the hour-long timescale of its biological function. The Ground state is dramatically destabilized at cold temperatures, and the system contains an alternate “off” conformation that exchanges with the primary “off” conformation at faster timescales than the rate-limiting step. Our findings demonstrate a simple principle for evolving a protein switch: one part of a protein domain remains stably folded to serve as a scaffold for the rest of the protein to re-fold.

## Introduction

The ability of biomolecules to sample multiple discrete conformations equips them with the capacity to sense and respond to arbitrary stimuli in their surroundings^1–3^. This fundamental principle was first laid out by Monod and co-workers in tandem with delineating the concept of conformational selection vs. induced fit.^4^ Their conclusions were derived from studying multimeric enzymes, though they raised the question, “do allosteric transitions occur in monomeric proteins containing a single polypeptide chain?”^1^ Indeed, monomeric allosteric behavior has since been demonstrated in systems ranging from signaling proteins^5^ to enzymes^6^. In more recent years, the discovery of “metamorphic” proteins have provided extreme examples of proteins switching their entire secondary structure between more than one conformational state^7^. The same questions first posed 60 years ago still apply -- for each of these discovered conformational changes, do they arise from conformational selection or induced fit? What allostery might be intrinsic to a single monomeric domain? What is the pathway of interconversion?

Several known metamorphic proteins have been demonstrated to spontaneously and reversibly interconvert between multiple states. Wild-type Mad2 regulates human cell cycle checkpoints, and interconverts spontaneously between monomer and dimer^8^. Lymphotactin, a chemokine, interconverts between a monomeric and dimeric form on the millisecond timescale^9^. Stopped-flow measurements as well as HDX-MS suggest that the mechanism proceeds through global unfolding^10^, though a more recent simulation study has suggested the presence of transiently folded intermediates^11^. IscU, an iron-sulfur cluster assembly scaffold protein, spontaneously interconverts between a structured and a primarily-disordered state that are both functionally relevant^12^.

KaiB is a circadian rhythm protein first studied in cyanobacteria in conjunction with KaiC, KaiA, and other partners^13–15^. In the studied variants from cyanobacteria, KaiB interconverts between a tetrameric Ground state and a monomeric “Fold-switched” (FS) state, which is the binding-competent state for KaiC and other binding partners. However, KaiB has since also been demonstrated to regulate circadian rhythms in bacteria lacking KaiA^16^. The KaiB from one of these bacteria*, Rhodobacter sphaeroides,* spontaneously interconverts between two monomeric forms.^16,17^, in striking contrast to KaiB from cyanobacteria.

Given the sparsity of knowledge for metamorphic proteins on possible mechanisms of interconversion between multiple stable conformations, how they evolved to occupy multiple states, and how these multiple conformations and the timing of their interconversion relates to function, we set out to characterize the conformational landscape of KaiB from *Rhodobacter sphaeroides* using real-time and relaxation NMR: Chemical Exchange Saturation Transfer (CEST)^18^ and Carr-Purcell Meiboom-Gill (CPMG)^19,20^ relaxation dispersion. By combining these three techniques, we can monitor all processes on the order of milliseconds to hours, and do indeed identify processes at all of these timescales.

We find that the rate-limiting step of KaiB switching from its binding-incompetent Ground state to the KaiC-binding competent FS state is the *cis-trans* isomerization of three prolines in the C-terminal domain, since the prolyl cis/trans isomerase CypA catalyzes the interconversion. The transition proceeds via a “partially-disordered” (PD) state, where the C-terminus is disordered but the N-terminus remains ordered. We further discover that KaiB is subject to cold denaturation as at 4°C, the PD state becomes the major state. Finally, we realize that KaiB interconverts with a fourth previously uncharacterized state which we term the “Enigma state”. We provide evidence from NMR chemical shift predictors that this fourth state is a register-shifted version of the Ground state as predicted by AF-Cluster^17^ and other adaptations of AlphaFold2^21^. Our work underscores the importance of experimental characterization of conformational landscapes, and that even for KaiB, which is one of the better-known fold-switching proteins, multiple surprises can await when systematically characterizing its mechanism of fold-switching.

## Results

### Interconversion between ground state and fold-switched state of KaiB takes hours

KaiB from *Rhodobacter sphaeroides* (KaiB^RS^) is a 92-residue protein that interacts with KaiC to regulate the circadian rhythm^16^. As part of this function, KaiB interconverts between a major, “Ground” state and a minor, “Fold-switched” (FS) state in solution^17^, where the FS state is the binding-competent state for KaiC (**Fig. 1a**, KaiB:KaiC structure: 8FWJ^16^). The N-terminal portion of both conformations (residues Met1-Glu51) have the same secondary structure (in grey, **Fig. 1a**), whereas the C-terminal domain switches secondary structure elements: in the Ground state, residues His52-Glu90 exist in a helix-helix-strand conformation, whereas in the FS state, the same stretch adopts a strand-strand-helix conformation. Unlike KaiB from *Synechococcus elongatus*, whose ground state exists primarily as a tetramer^13^, the Ground state of KaiB^RS^ exists as a monomer. In addition, the wild-type protein already populates the FS sufficiently to be directly observable in NMR experiments, making KaiB^RS^ (henceforth KaiB) an attractive system to investigate the mechanism of how the protein interconverts between these two very different folded states.

**Figure 1.**
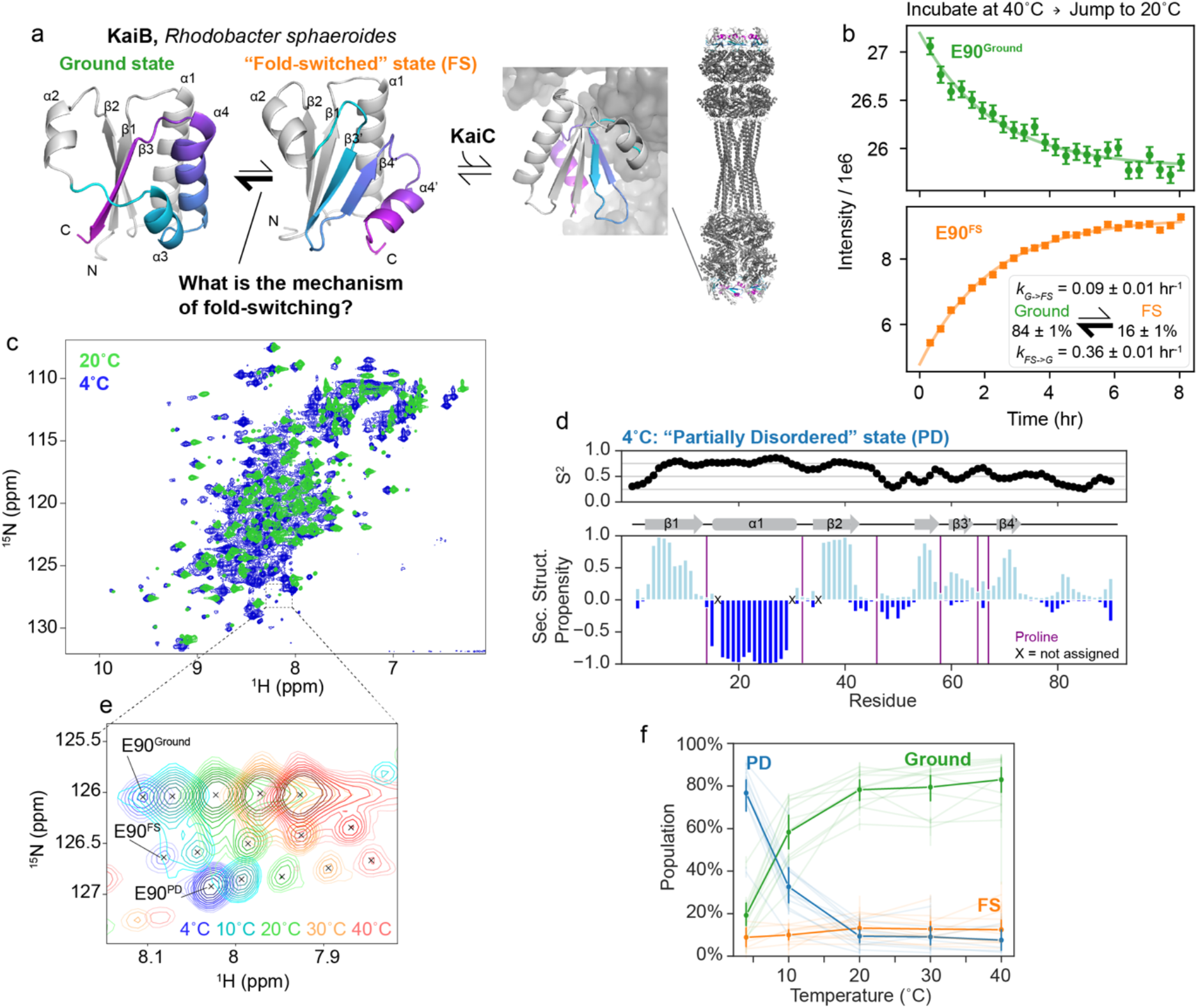
KaiB interconversion between ground and FS state on the hour time regime. a) Current understanding of the KaiB:KaiC system in *Rhodobacter sphaeroides.* Monomeric Ground state and fold-switched “FS” state models were generated with AF-Cluster^17^ with the fold-switching part colored for the same sequence regions. Structure of KaiB:KaiC complex: 8FWJ^16^. b) Perturbing populations of the Ground and FS state by incubating at 40°C and jumping to 20°C followed by real-time NMR spectra to quantify the interconversion rates at 20°C depicted in the inset. c) Comparing ^15^N-HSQC spectra at 4°C and 20°C reveals peaks in the disordered region of the spectrum at 4°C not present at 20°C. d) Predicted backbone order parameters and secondary structure propensity of the major state at 4°C. The N-terminus retains the same fold as found in the Ground and FS states at 20°C, but the C-terminal domain is more disordered. e) Representative temperature-dependent peak shifts and population changes of the three states. f) Populations for eleven residues for which the Ground, FS, and PD states could be assigned in the majority of temperature conditions, calculated from peak volume. The solid lines represent average populations at each temperature.

Our first question was, what is the rate of interconversion between the Ground and FS state? Given that KaiB is involved in regulating circadian rhythms, a biochemical cycle which takes hours, we were curious if the interconversion rate was similar to the slow timescale of the circadian rhythm^16^. If so, we reasoned that we could use NMR in real-time to monitor the rate of interconversion between states by perturbing the populations with a temperature-jump experiment. By incubating at 40°C, then jumping to at 20°C and collecting consecutive ^15^N-HSQC spectra, we indeed monitored interconversion dynamics between the Ground and FS state, quantified via peak intensities (**Fig. 1b**). We calculated interconversion rates of *k_G→FS_* = 0.09 ± 0.01 hr^-1^ and *k_FS→G_* = 0.36 ± 0.01 hr^-1^ from a global fit of 45 Ground state peaks and 37 FS state peaks (**Fig. 1b, inset**). The timescale of Ground ↔ FS interconversion is indeed slow: to put another way, the mean lifetime of the Ground state, 1/*k_G→FS_,* is 11 ± 1 hrs, and the mean lifetime of the FS state, 1/*k_FS→G_,* is 2.8 ± 0.1 hrs.

### KaiB partially denatures at cold temperatures

In ^15^N-HSQC spectra of KaiB, we noticed a small population of a third conformation that could not be assigned to either the Ground or FS state. This population strongly increased with lowering the temperature to the point that at 4°C this conformation became the dominant species (**Fig. 1c**). We were therefore able to characterize the structural identity of this third, unknown conformation. It appeared to predominantly occupy a disordered state, first evidenced by peaks observed in the center of a ^15^N-HSQC spectrum (**Fig. 1c**). After assigning the backbone of this major state at 4°C, we predicted backbone order parameters (S^2^) and secondary structure propensities using RCI^22^ and TALOS-N^23^, respectively, (**Fig. 1d**) to better understand the structure of this major state populated at 4°C. The N-terminus of KaiB (1-48) was predicted to retain high S^2^ as well as the same secondary structure as the Ground and FS states. In contrast, the C-terminal part – the “switch region” – has low S^2^ values and little secondary structure propensity, indicating a disordered region. Furthermore, the backbone chemical shifts of the C-terminus are in excellent agreement with chemical shifts for random coil as predicted with POTENCI^24^, with a net RMSD of 0.36 ppm in the C-terminal region over N, H, C𝘢, and Cβ atoms (**Fig. S1**). We thus designated this new state the “partially disordered” (PD) state. Intriguingly, this “PD” state is indicated to have more beta-stranded character, in particular in residues 54-72, suggesting transient pre-forming of the beta-stranded FS state (**Fig. 1d**).

We recorded ^15^N-HSQC spectra at temperatures ranging from 4°C to 40°C to monitor the temperature dependence of the Ground, FS, and PD states. Representative temperature shifts of the C-terminal peak are depicted in **Fig. 1e**. We estimated the populations of the three states via peak intensities for well-resolved cross peaks for which all states could be assigned (**Fig. 1f**). At 4°C, the PD state was populated to about 75%, which dropped to 30% at 10°C. As temperature increases, the Ground state becomes increasingly more populated, reaching roughly 80% occupancy by 20°C. The population of the FS state remains between 10-20% across the temperature range.

To test if KaiB’s partial denaturation at 4°C was reversible, we incubated KaiB at 4°C and then monitored populations using real-time NMR at 20°C, analogously to our previous temperature-jump experiment (**Fig. 2a**). We observed that KaiB did indeed return to Ground and FS populations identical to those previously quantified at 20°C (**Fig. S2**). Intriguingly, although the PD state was populated at roughly 75% at 4°C, the PD state population was about 10% at the first collected HSQC after 7 minutes of deadtime and remained constant over the course of the experiment (**Fig. 2a**). This indicates that interconversion in and out of the PD state is significantly faster than the interconversion between the Ground and FS state, but the connection between the three states remained undetermined. Next, we set out to determine if this “partially disordered” (PD) state represented an on-pathway intermediate for the interconversion between the Ground and FS states, or a separate off-pathway process.

**Figure 2.**
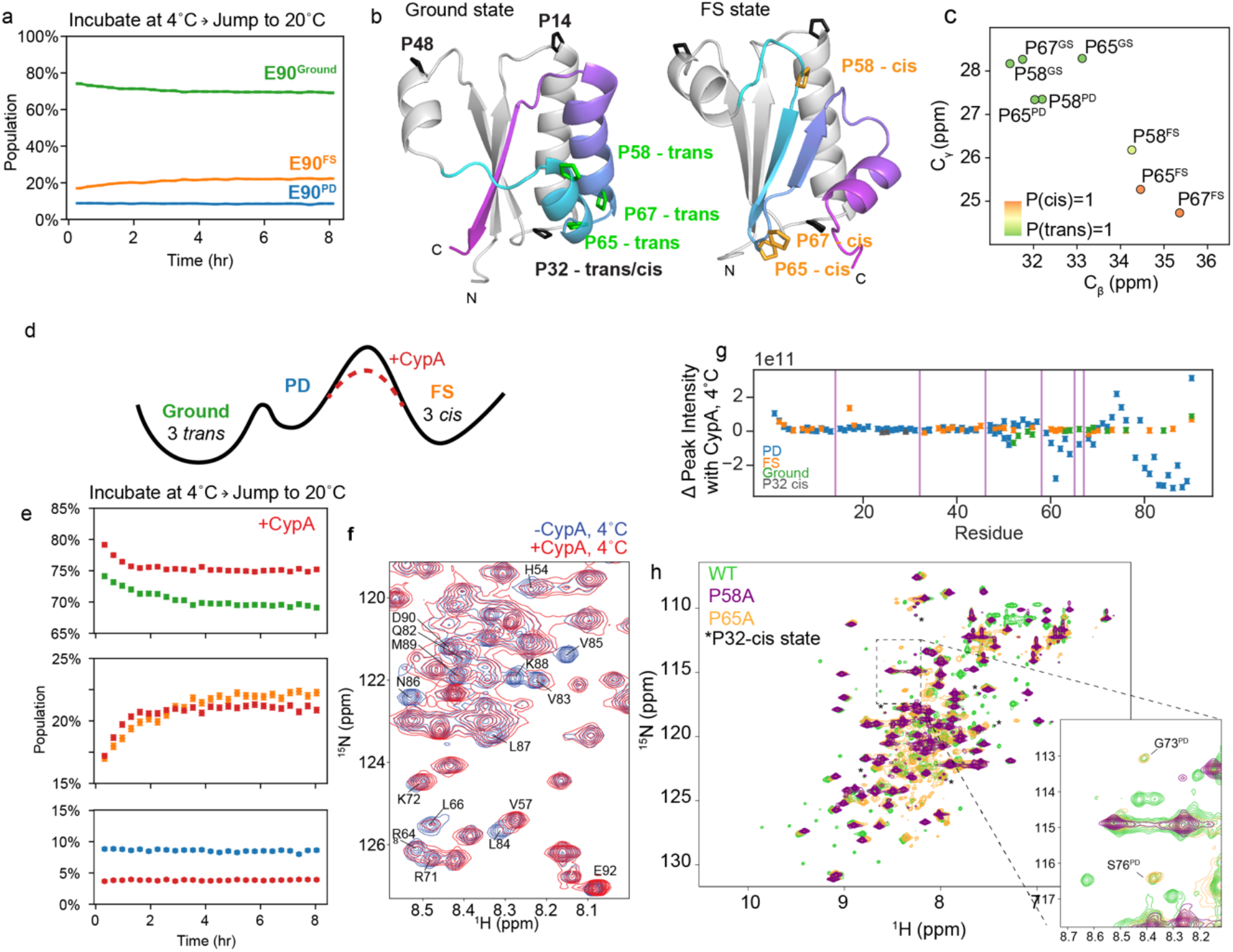
KaiB interconverts between the Ground and FS state via the partially disordered state, and proline cis/trans isomerization is rate-limiting. a) Incubating at 4°C and jumping to 20°C followed by ^15^N-HSQC time series at 20°C reveals that KaiB cold denaturation is reversible, and that interconversion in and out of the PD state at 20°C is significantly faster than the deadtime of this experiment (7 min). b) Visualizing location of prolines and their difference between *trans* and *cis* conformations in Ground and FS states. Monomeric Ground state and fold-switched “FS” state models were generated with AF-Cluster^17^. c) C_β_ and Cᵧ chemical shifts of P58, P65, and P67 in Ground, PD, and FS state, colored by cis/trans probabilities predicted with PROMEGA^25^. d) Scheme of proposed mechanism for accelerating interconversion by adding the peptidyl prolyl cis/trans isomerase Cyclophilin A (CypA). e) Adding 30 µM CypA to the temperature-jump experiment of 1.2 mM KaiB accelerates the Ground to FS interconversion rates by 2.5-fold. f) ^15^N-HSQC spectra indicate peak broadening for residues in the C-terminus of the partially disordered state upon addition of CypA. g) Change in peak intensities across all assigned states at 4°C indicates that CypA binds to the C-terminus of the PD state. h) Single point mutations of prolines P58 or P68 are sufficient to destabilize the FS state.

### Triple-Proline isomerization is responsible for slowness of Ground-FS interconversion

In speculating on why the interconversion between Ground and FS state takes several hours, we noticed that there are three proline residues in the fold-switching C-terminus (P58, P65, and P67) that occupy a *trans* conformation in the Ground state and a *cis* conformation in the FS state (represented using AF2 models in **Fig. 2b**). This is supported by C_β_ and Cᵧ chemical shifts (**Fig. 2c**) and homologous structures for both states from other organisms (**Table S1**). In the PD state, P58 and P65 have C_β_ and Cᵧ chemical shifts corresponding to the *trans* state (**Fig. 2c**). P67 in the PD state Cᵧ was unable to be identified in the CCONH experiment used to assign Cᵧ’s as it was collected at 20 °C where the PD state is minorly populated. Using the P67 Cβ chemical shift (32.22 ppm) and neighboring chemical shifts, the PROMEGA^25^ algorithm predicts P(trans)=0.90. The evidence that these three prolines are primarily in *trans* in the PD state matches our expectations: a peptidyl-prolyl bond in a random coil is favored to be in *trans* ^26,27^.

We hypothesized that the Ground state can interconvert relatively fast with the PD state maintaining these three prolines primarily in trans, and that the *trans→cis* interconversion of the three prolines is rate-limiting for the conformational change to the FS state. How can we experimentally test such a mechanism? Cyclophilin A (CypA), a peptidyl-prolyl cis/trans isomerase, can catalyze this reaction provided that the X-Pro bonds are accessible, as would be the case in the PD state (scheme in **Fig. 2d**). Indeed, adding 30 µM CypA to 600 µM ^15^N-labeled KaiB immediately prior to recording the temperature-jump experiments increased both the forward and reverse rates roughly 2.5-fold (example traces in **Fig. 2e**), to *k_G→FS_* = 0.24 ± 0.02 hr^-1^ and *k_FS→G_* = 0.98 ± 0.04 hr^-1^, buttressing our hypothesis that the cis/trans isomerization of these three prolines is the rate-limiting step for the fold switching.

To further validate that CypA does indeed bind the PD state, we collected ^15^N-HSQC spectra at 4°C, where the PD state is most populated, to monitor peak intensity changes upon addition of excess CypA. We indeed observed cross peaks corresponding to the PD state were shifted and/or missing in the presence of CypA (**Fig. 2f**). **Fig. 2g** depicts calculated changes in cross peak intensities upon addition of CypA for all states assignable at 4°C. Significant changes are observed from roughly residue 50 onwards in the PD state, and not in the Ground or FS states, supporting our hypothesis that CypA binds the PD state of KaiB, and that therefore the rate-limiting step for Ground-FS state interconversion is *cis-trans* isomerization in the PD state. Underscoring their importance, these three prolines are conserved across 474 of 487 KaiB variants (97%) in the KaiB phylogenetic tree in ref. ^17^ (**Fig. S3**).

An additional test for the importance of the prolyl cis/trans isomerization for the slow fold switch in KaiB was to mutate any of these prolines. Mutating already a single proline, either P58 or P65 to alanine resulted in a complete loss of peaks corresponding to the FS state (**Fig. 2h**), strengthening the proposed mechanism in which all three prolines need to switch to *cis* to populate the FS from the PD state. Peaks for G73 and S76 in the PD state were visible for the P65A mutant (inset, **Fig. 2h**). There were some minor peaks still present in spectra of the P58A and P65A mutants, identified with asterisks in **Fig. 2h**, which aided us in identifying that P32, in a loop of the N-terminus, undergoes a cis-trans isomerization independently of the Ground ↔ PD ↔ FS conformational change (**Fig. S4**).

### The Ground state of KaiB interconverts with a 4^th^ “enigma” state

Given our finding that the Ground state interconverts to the FS state via a partially-disordered state, we wanted to better characterize the kinetics of this conformational exchange process that were too fast to monitor in the temperature-jump experiments. We turned to chemical-exchange saturation transfer (CEST) NMR^18^, which allows for monitoring millisecond-timescale processes, delivering the populations, rates, and chemical shifts of the interconverting species. At both 4°C and 20°C, we saw signs of several processes in our CEST data. **Fig. 3a** shows CEST traces at 4°C for a representative residue Asn84 (data for all fit residues in **Fig. S5**). The Ground state trace has two dips, indicating interconversion with two other states (**Fig. 3a, top**). One of these states (at B_1_ = 122.3 ppm, blue line) is indeed the PD state, and exchange with it is observed as reciprocated when monitoring the PD state (**Fig. 3a, bottom**). However, there is another state (B_1_ = 125.2 ppm, purple line) also observed to interconvert only with the Ground, not with the PD state. At 20°C, the PD dip decreases in size (in agreement with the small population of PD at that temperature), but there is still clearly exchange of the Ground state with the new state at 125.2 ppm, depicted in purple in **Fig. 3a,b**. Notably, this is *not* the FS state, which is assigned at 20°C at 117 ppm (**Fig. 3b**, bottom) and does not have any observable CEST exchange. Not knowing the structure of this fourth state, we designated it the “enigma” state.

**Figure 3.**
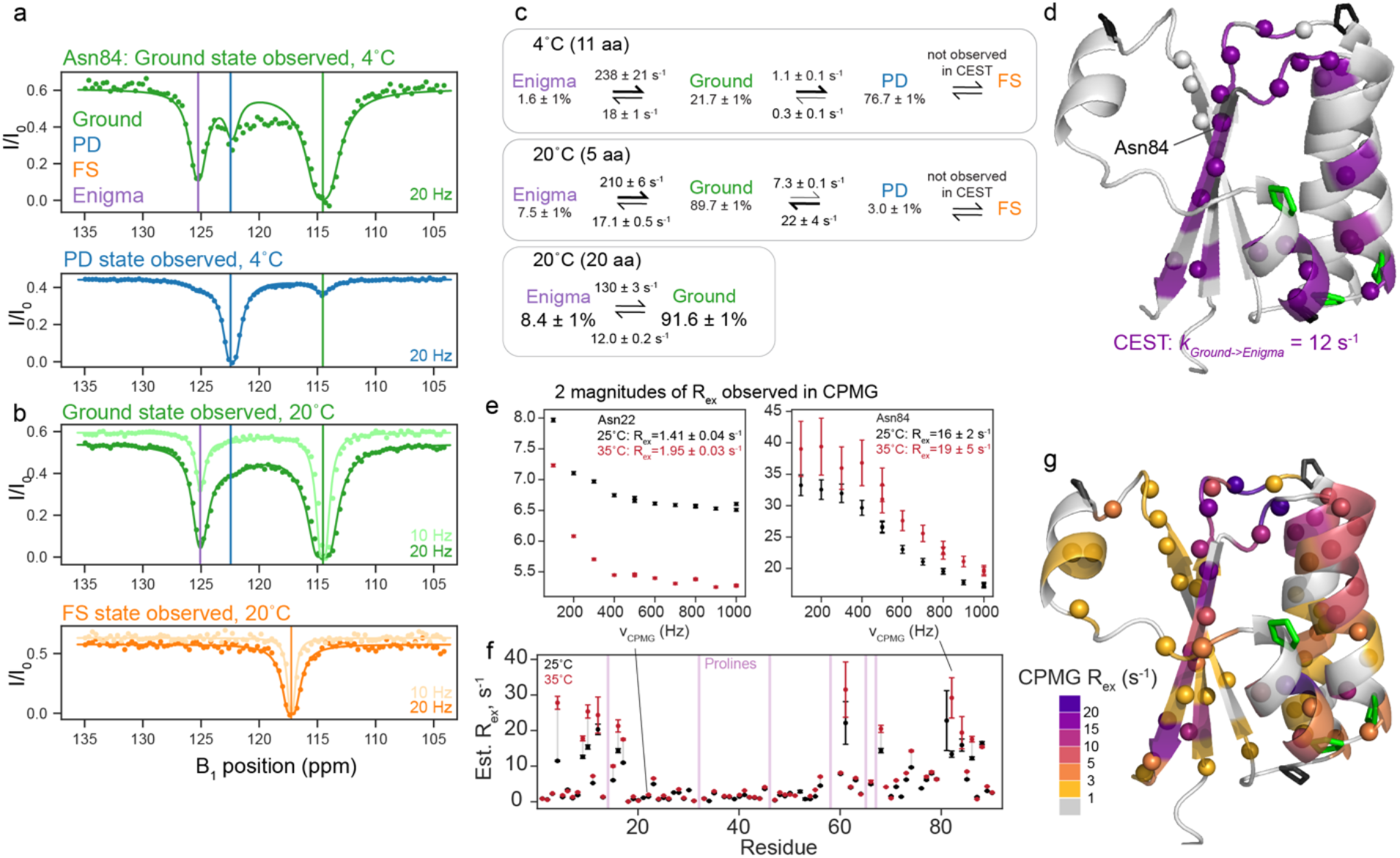
The KaiB Ground state interconverts with a fourth “enigma” state. a) Representative ^15^N backbone CEST data at 4°C for Asn84, indicating that the Ground state exchanges with both the PD state and another state. b) CEST data at 20°C for Asn84, indicating that at 20°C, exchange with the Enigma state is still present, and it is not the FS state. c) Populations and interconversion rates for models of exchange between the Enigma, Ground, and PD states determined by global fits of the number of residues indicated (see also Figs. S5 and S6). d) All residues with signs of exchange at 20°C are indicated with spheres. Residues fit to one two-state process (Ground to Enigma state) are colored in purple. e) Representative ^15^N backbone CPMG data of residues with small R_ex_ values (Asn22) and larger R_ex_ values (Asn84). f) Estimated R_ex_ values from CPMG at 25°C and 35°C. The majority of residues have greater R_ex_ with increasing temperature, indicating the CPMG is reporting on slow-intermediate-timescale processes that is consistent with the processes observed by CEST. g) Estimated R_ex_ values for all residues with R_ex_ > 1 s^-1^ visualized on the structure. The residues with CPMG R_ex_ > 10 s^-1^ agree with the residues exchanging with the Enigma state, reported by CEST. Residues with very small R_ex_ are possibly reporting on a putative cis-trans isomerization (see Figures S4 and S8).

The minimal model to fit our data at both temperatures follows the scheme Enigma ↔ Ground state ↔ PD state (**Fig. 3c**). The enigma state is populated to about 8% at 20°C and 2% at 4°C. At both temperatures, *k_G→PD_* is significantly slower than *k_G→E_*. 20 residues showed two-state exchange at 20°C to peaks which were not the chemical shift of the PD state, and were therefore the Enigma state. We used these residues to fit a two-state model to better understand the Ground↔ Enigma process. The locations of all 20 residues with exchange to the Enigma state are depicted on the Ground state structure in **Fig. 3d**. This group includes most residues in the C-terminus as well as residues 12-14. Details of all fits compared are in Methods and **Fig. S6**.

To assess if KaiB contained any processes too fast to detect via CEST, we also collected CPMG relaxation dispersion^19,20^ data at 25°C and 35°C (representative data in **Fig. 3e**, all data in **Fig. S7**). Residues with R_ex_ fell into two general categories: residues with R_ex_ > 10 s^-1^, and residues with R_ex_ of about 2 s^-1^. Most residues in both classes had increased R_ex_ with increasing temperature (**Fig. 3f**), indicating both processes are on the slow to intermediate timescale (see Methods), and that the R_ex_ measured with CPMG is estimating the rate of leaving the observed state^28,29^. Note this means that the CPMG data could not be further analyzed with the Carver-Richards equation, but provided qualitative support for our results obtained from CEST. **Fig. 3g** depicts the magnitude of R_ex_ calculated at 25°C for residues with R_ex_ > 1 s^-1^ colored on the Ground state structure. Residues that were fit via CEST to exchange with the Enigma state as one global process (purple in **Fig. 3d**, compare to purple in **Fig. 3g**) also predominantly had R_ex_ ∼ 12 s^-1^ measured by CPMG, which suggests that both CEST and CPMG are reporting on the same process of Ground -> Enigma state, where *k_G->E_* is roughly 12 s^-1^.

Many other residues throughout KaiB were observed to have a R_ex_ of roughly 2 s^-1^. We postulate these are reporting on the *k_G->PD_* process, as the rate fit via CEST matches the R_ex_ from CPMG, but some may also be reporting on a separate process: residues 39-42 at the top of strand β 1 could be fit to with *k_ex_* = 6.4 s^-1^ and an alternate-state population of 5% (**Fig. S8**). In conclusion, by collecting CPMG at two temperatures, we ascertained that the processes observed were in the slow-intermediate regime, in agreement with our CEST data, and that there were no processes at a faster timescale detectable.

### AlphaFold and NMR chemical shift predictors suggest the Enigma state is a β strand register shift

What is the structure of this new enigma state? The structure model that we have used to visualize the Ground state of KaiB is the one returned with the highest pLDDT from AF-Cluster^17^ (**Fig. 4a**). This structure aligns to the crystal structure of KaiB from *T. elongatus*^30^ with 0.69 Å RMSD, and for clarity we henceforth denote as the “TE-like” structure. However, we realized that among the top 5 highest-pLDDT Ground state models in our AF-cluster analysis^17^, some contained a subtle difference: a register shift of 2 amino acids in the hydrogen bonding between the β1 and β3 strands (**Fig. 4b**). The register-shifted state has a different, repacked hydrophobic core (**Fig. S9**). Predicted free energies from FoldX^31^ (**Table S2**) indicate that the TE-like state has a more favored hydrophobic core, yet the register-shifted state allows four charged residues in the fold-switching region to be more solvent-exposed than they are in the TE-like state. Which of these two structure models – the TE-like structure, or the register-shifted structure – corresponds to the major Ground state conformation for KaiB^RS^? We collected ^15^N-edited NOESY data and compared both sets of beta-sheet hydrogen bonding to it, and found that the higher-pLDDT TE-like model (“TE-like”) matched the major observed hydrogen-bonding state (**Fig. S10**). Could it be possible that the other structure predicted by AF-Cluster is the Enigma state?

**Figure 4.**
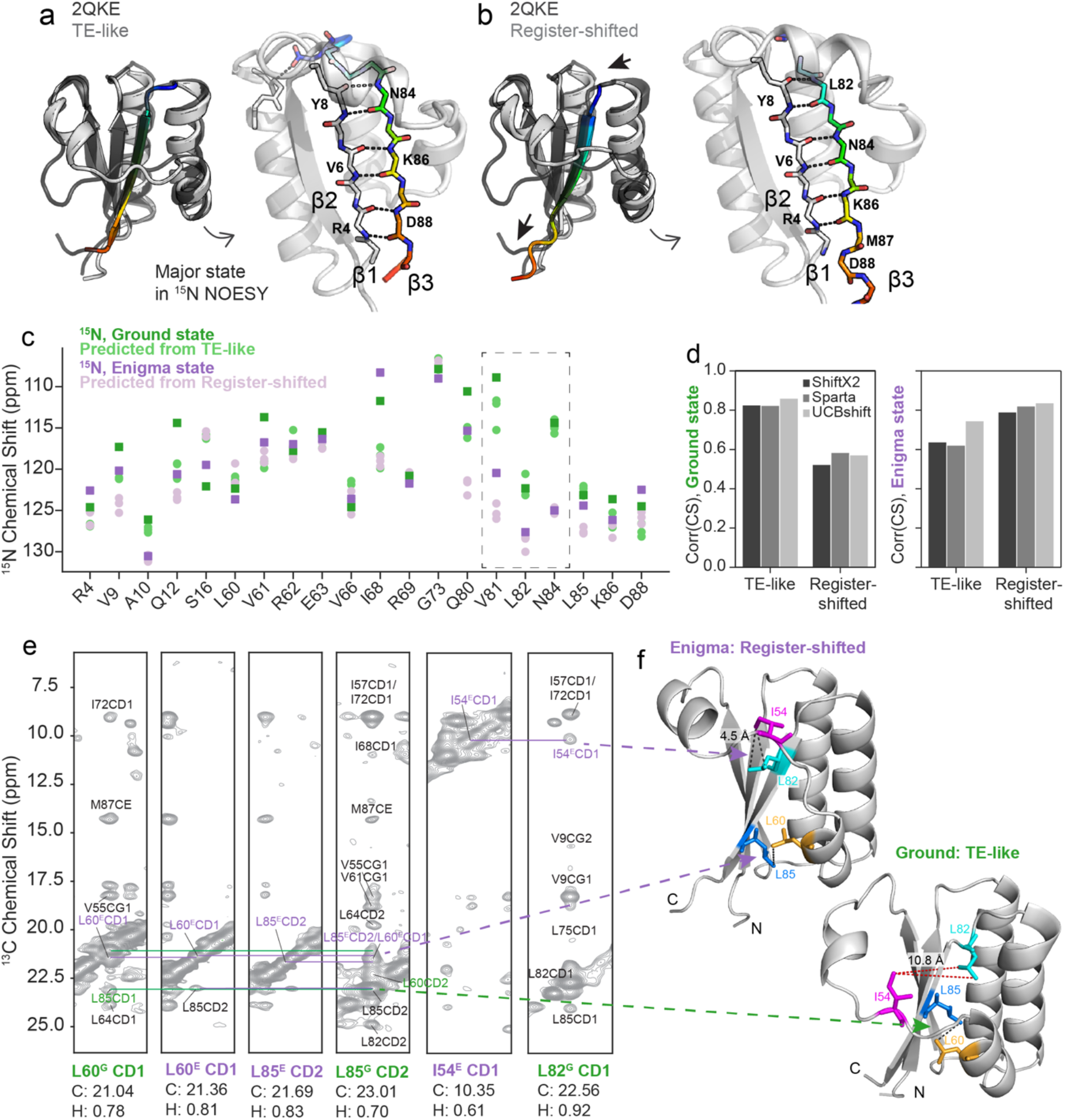
A subtle β1-β3 register shift predicted by AlphaFold2 and experimentally supported by NMR is consistent with the Enigma state. a) The highest-pLDDT KaiB ground state produced by AF-Cluster^17^ has the same hydrogen-bonding as KaiB from *Thermosynechococcus elongatus* (PDB: 2QKE^30^). We refer to this state in KaiB^RS^ as “TE-like”. Left: superposition of TE-like state and 2QKE. Right: detail of beta-strand hydrogen bonding. b) AF-Cluster, and AF2^21^ with single-sequence sampling, also predict another ground-like state with lower pLDDT where β3 is register-shifted by 2 amino acids. Arrows indicate direction of register shift. Left: superposition of register-shifted state and 2QKE, right: detail of altered beta-strand hydrogen bonding. c) Visual comparison of ^15^N chemical shifts for the ground state (green) and enigma state (purple) with chemical shift predictions from 3 chemical shift predictors (SHIFTX2^33^, SPARTA+^32^, and UCBshift^34^) using the TE-like model (light green) and the register-shifted structure (light purple). d) Comparing correlations between predicted and measured chemical shifts for 3 tested chemical shift predictors shows that predictions from the TE-like structure correlate better with the Ground state, and predictions from the Register-shifted structure correlate better with the Enigma state. e) Methyl-methyl NOESY supporting contacts unique to the register-shifted structure, contacts depicted in (f). In the register-shifted structure, Leu82 is on the exterior of the b3 strand and can interact with Ile54, but in the TE-like structure, it is flipped into the interior to form the hydrophobic core (right), and is not within NOE distance of Ile54. Also in support of the register-shifted structure is two distinct sets of cross-peaks between L60 and L85, one corresponding to Ground state chemical shifts and the other to Enigma methyl chemical shifts.

We were curious if the sparse experimental data we had in hand on the Enigma state – namely, 20 ^15^N chemical shift values – in combination with predicted ^15^N chemical shifts from these structure models, could offer any insight. We predicted chemical shifts for the 2 different structural models using three independent algorithms: SPARTA+^32^, SHIFTX2^33^, and UCBshift^34^. The predicted values and the experimentally determined ^15^N shifts of the Ground and Enigma states are compared in **Fig. 4c**. We were surprised to find striking agreement between the experimental Enigma chemical shifts and the predicted chemical shifts from the register-shifted model, particularly for the residues with the largest changes in chemical shift between the Ground and Enigma state: Val81, Leu82, Asn84, values boxed in **Fig. 4c**. Val81 and Leu82 dramatically switch places between the Ground state and register-shifted structure – in the TE-like structure, Val81’s sidechain is in the core and Leu82 is exposed, and in the register-shifted structure, Leu82’s sidechain participates in the core and Val81 is facing the surface (**Fig. S9**). Across all three algorithms, chemical shifts predicted from the TE-like structure correlated best with the experimental Ground state chemical shifts, and conversely, chemical shifts predicted from the Register-shifted structure correlated best with experimental Enigma state chemical shifts (**Fig. 4d**).

How could we experimentally determine if this structure is present? We could not identify any cross peaks in ^15^N HSQC corresponding to the Enigma state chemical shifts observed in CEST, but hypothesized that methyl NMR might be more sensitive to this minor state that is only populated to 8%. Indeed, we found that by combining chemical shift information from methyl CEST^35^ and CPMG^36^ (see Methods, **Figs. S11**-**S13**), we could assign a few resonances corresponding to the Enigma state and identify NOE cross-peaks that are due to exchange between the enigma and ground states (**Fig. 4e**). Notably, we found a contact uniquely present in the register-shifted state between Ile54 CD and Leu82 CD. This forms part of the repacked hydrophobic core of the register-shifted structure compared to the TE-like structure. In the TE-like structure, Leu82 forms the hydrophobic core (**Fig. 4f**, bottom), but in the register-shifted structure, has flipped to the exterior, and is therefore within NOE distance of Ile54 (**Fig. 4f**, top). Furthermore, we found two distinct sets of NOEs between L60 and L85, one corresponding to Ground state chemical shifts and the other to Enigma methyl chemical shifts. The presence of these NOEs suggests that the register-shifted structure is indeed the Enigma state, and this register-shifted β1-β3 conformation was further validated by unique NOE’s in ^15^N and ^13^C edited 3D NOESY (**Fig. S10**).

To assess if we could identify any structural models with even higher correlation to the Enigma state, we predicted chemical shifts in UCBshift for all models produced by AF-Cluster^17^, as well as eighty models sampled from AlphaFold2 (AF2) in single-sequence mode^21^ (**Fig. S14**). We selected UCBshift^34^ as it was the algorithm that achieved the highest correlation to the Enigma state by a moderate amount. AF2 in single-sequence mode predicted only the Enigma state, indicating in retrospect that AF-Cluster was required to generate a model that best matches our experimental data of the Ground state. We did not find any structure models from these sampling techniques that resulted in chemical shift predictions higher correlation to the experimental Enigma state chemical shifts.

### Free energy landscape of KaiB dynamics

Figure 5a summarizes the structures, populations, and timescales of the dynamic processes of KaiB delineated in this work. The rate-limiting step for KaiB to interconvert between the Ground and FS state is the cis-trans isomerization of three prolines in the C-terminus. This proceeds via a “partially disordered” state that is populated to 3% at 20 °C, but the major state at 4°C. In this state, residues up to Glu51 in the N-terminus retain the same secondary structure as in the other folded states, whereas C-terminal residues are disordered. By using CEST to characterize the dynamics of interconversion between the Ground and PD state, we discovered that the Ground state also interconverts with a 4^th^ “enigma” state on the millisecond timescale, which we postulate is a register shift in the hydrogen bonding between the β1 and β3 strands.

**Figure 5.**
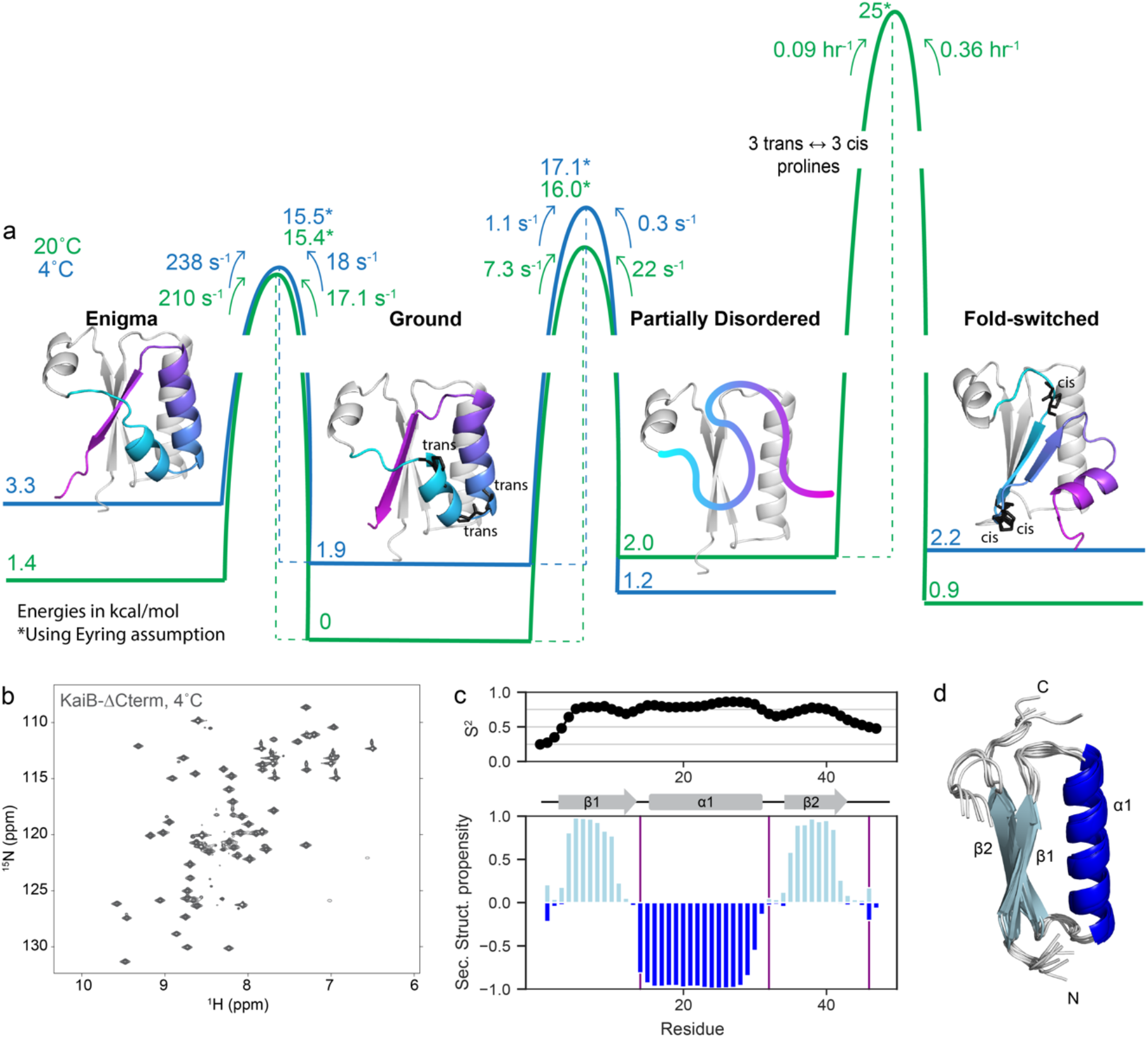
The conformational landscape of KaiB is unified by a stable subdomain. a) Calculated free energies of all states and barrier heights, incorporating information from temperature-jump and CEST experiments. Models for the enigma, ground and fold-switched state were generated with AF-Cluster^17^. The model for the partially disordered state is the N-terminus spliced from the Ground state model. b) The proposed mechanism of fold-switching via a partially-disordered intermediate suggests that the N-terminus of KaiB should be stable as an independent unit. Indeed, KaiB-ΔCterm (KaiB^1–46^) is stably folded at 4°C as shown via ^15^N-HSQC. c) Predicted backbone order parameter S^2^ and secondary structure propensity for KaiB-ΔCterm. d) Calculated structure of KaiB-ΔCterm using CS-Rosetta^43^.

In all these states, the N-terminus remains constant, and suggests that throughout the fold switch, the N-terminus remains stably folded as a single unit. To test this, we produced a truncation of KaiB terminating at Leu46 (“KaiB-ΔCterm”). Strikingly, KaiB-ΔCterm indeed remained stably folded over the entire temperature range (Fig. 5b and **Fig. S15**), and NMR C𝘢 and Cβ assignments collected at 20°C exhibited predicted secondary structure propensities identical to the corresponding N-terminal region of full-length KaiB (Fig. 5c; *cf.* Fig. 1d, 25°C ^15^N-HSQC in **Fig. S15**). A structure prediction with CS-Rosetta supports our expected model for this miniprotein (Fig. 5d).

## Discussion

Some members of the KaiB family do not fold-switch, and are stabilized in the FS state^17,37^. We postulate that fold-switching KaiB variants likely evolved from variants that only occupy the FS state, as the thioredoxin-like fold of the FS state is an ancient, very common fold, contrary to the Ground state fold, which has been identified only in the KaiB family^13^. Notably, P58 in KaiB is a proline that is conserved, and in the *cis* conformation in several redox protein families with thioredoxin-like folds^38^ and thought to be essential for redox function by forming part of the binding site for substrates^39^. The prolines at positions 65 and 67 likely evolved later, adding the capability to finally toggle with an inactive form for its use in circadian rhythm.

Our finding that the N-terminus of KaiB can independently fold in a stable conformation hints at a mechanism for how KaiB could have evolved: a gene for a protein with a thioredoxin-like fold that performs a different biological function is duplicated to create a proto-KaiB that still only occupies only the FS state. To toggle the binding of this proto-KaiB to KaiC and other circadian rhythm proteins, the C-terminus could incur point mutations that cause it to also occupy a disordered state to disrupt the binding surface in this alternate conformation. However, a partially-disordered protein could cause nonspecific binding or aggregation, which would make the evolution of a second folded state evolutionarily advantageous. The actual nature of this second state – i.e., the Ground state – perhaps matters less, only that it is kinetically separated from the FS state at a hours-long timescale for modulating circadian rhythms. This hypothesis is supported by the fact that the ground state interconverts with the Enigma state in our variant, while other variants tetramerize. Evidently, evolution can be allowed flexibility in designing the “off-state” of a switch.

There are likely many more details and general principles to be understood in how proteins switch folds from one to the other and how this is related to their biological function. This is evidenced by the number of open questions about the interconversion pathways of other metamorphic proteins. Lymphotactin is a metamorphic protein that interconverts between a monomeric Ltn10 fold and a dimeric Ltn40 fold. Its mechanism of interconversion remains controversial; stopped-flow and fluorescence measurements have been interpreted to mean that lymphotactin completely unfolds^10^, yet more recent molecular dynamics experiments suggest that there could be partially-ordered intermediates that the pathway proceeds through^11^. RfaH is a metamorphic protein with a C-terminal domain that interconverts between an alpha-helical and beta-stranded form. Molecular dynamics methods using different sampling methods to understand the interconversion mechanism of the metamorphic protein RfaH have resulted in different proposed interconversion pathways.^40^

The study of the dynamics and interconversions of single proteins is already nontrivial, and a complicating factor in understanding how a given interconversion pathway might have evolved is the myriad other biomolecular interactions possibly relevant. This is exemplified by the metamorphic protein Mad-2, which interconverts between two distinct folds^8^. Its interconversion is linked to binding Cdc20 through the so-called “seatbelt” mechanism, which is rate-limiting for the assembly of the mitotic checkpoint complex (MCC)^41^. Absent of any other binding partners, its interconversion takes hours, but MCC assembly in the cell is complete in minutes^42^. Numerous other binding partners accelerate the interconversion, though the exact biologically-relevant mechanism for fold-switching remains unknown. Some have used time-resolved NMR and relaxation measurements to dissect the mechanism of Mad-2 interconversion in the absence and presence of the Cdc20 MIM domain. However, the fold-switching speedup of this MIM domain alone is insufficient to account for assembly in the cell finishing in minutes^42^, and there are likely relevant interactions unaccounted for by these existing NMR studies. An open challenge for the field is how to detect transient populations present in these interconversion pathways in interaction with multiple other binding partners, ideally in the crowded interior of the cell.

In conclusion, we demonstrated that the slowness of a process that is necessitated by its biological function – namely, timekeeping for circadian rhythm regulation – can be encoded in the multiple conformations and interconversion mechanisms of a single protein domain. The diversity of interconversion timing even among known metamorphic proteins is astonishing. Lymphotactin interconverts on a timescale of seconds, perhaps essential for its role in immune signaling. Mad2 interconverts to complete its role in cell division in a matter of minutes. KaiB interconverts on a timescale of hours to facilitate the hours-long circadian rhythm. These are all examples of timing linked to function. We postulate that it is a general principle of protein conformational changes – that the timing of these conformational changes is subject to pressures of evolution, same as the conformations themselves.

## Acknowledgments

We thank Ricardo Padua, Hannes Ludwig, and other members of the Kern lab for fruitful discussion. This study made use of the National Magnetic Resonance Facility at Madison (NMRFAM), which is supported by NIH grant R24GM141526. We would also like to thank Dr. Marco Tonelli from NMRFAM for assistance with data collection. H.K.W.S. acknowledges funding from the Jane Coffin Childs foundation. This work was supported by the Howard Hughes Medical Institute (HHMI) to D.K.

## Competing Interests

D.K. is a co-founder of Relay Therapeutics and MOMA Therapeutics. The remaining authors declare no competing interests.

## Author Contributions

H.K.W.S., R.O., W.P., A.M.O., and D.K. designed the research. R.O., W.P., A.M.O. collected the NMR data. R.O., W.P., A.M.O., L.C. contributed to sample preparation and data collection. H.K.W.S., R.O., W.P., A.M.O., A.G. contributed to data analysis and interpretation. H.K.W.S., A.M.O., and D.K. wrote the manuscript.

## Methods

### Protein expression and purification

KaiB from *Rhodobacter sphaeroides* was expressed and purified as described previously ^17^. Wild-type Cyclophilin A (CypA) was expressed and purified as described previously^44^. In brief, plasmid encoding CypA was transformed into BL21(DE3) cells, plated on LB plates with ampicillin resistance, and grown overnight at 37°C. Single colonies were selected and grown overnight at 30°C. Cells were grown at 37°C to an OD_600_ ∼1.1 after which protein expression was induced with 0.3 mM IPTG for 4 h at the same temperature. Cell pellets were resuspended in lysis buffer containing 25 mM MES pH 6.1, 5 mM b-mercaptoethanol, 1x EDTA-free protease inhibitor cocktail (Thermo Fisher Scientific), DNase I (Sigma-Aldrich), lysozyme (Sigma-Aldrich). Lysate was sonicated on ice for 10 minutes (20s on, 40s off, output power of 35 W), followed by centrifugation at 18,000 rpm for 45 minutes at 4°C. Supernatant was purified on a SP-sepharose column using a NaCl gradient. Fractions containing CypA were pooled and dialyzed overnight at 4°C into 50 mM Na_2_HPO_4_, pH 6.8, 2 mM b-mercaptoethanol. Remaining impurities were removed using a Q-sepharose column. CypA was purified on a size-exclusion column (S75) equilibrated in 50 mM Na_2_HPO_4_, pH 6.7, 1mM TCEP, 250 mM NaCl, 5% glycerol. Prior to use, CypA samples were dialyzed overnight at 4°C into 100 mM MOPS at pH 6.5, 50 mM NaCl, 2 mM TCEP. For methyl dynamics experiments, a double methyl labeled ILVM (Ile, Leu, Val, Met) sample was prepared with [U-^15^N,^12^C,^2^H] {Ile (^13^C^δ1^H_3_), Leu(^13^CH_3_,^13^CH_3_), Val(^13^CH_3_,^13^CH_3_)}, Met(^13^CH_3_) labeling. The cells were grown at 37C in ∼99% D_2_O minimal M9 media containing 3 g/l [^2^H, ^12^C] glucose as the carbon source, and 1.5 g/l ^15^NH_4_Cl ammonium chloride. Methyl labeling was achieved by the addition of 75 mg/L of alpha-ketobutyric acid [U-^12^C, ^2^H, methyl-^13^CH3] for labeling the isoleucine, 100 mg/L of α-ketoisovaleric acid [U-^12^C, ^2^H, methyl-(^13^CH3, ^13^CH3)] for labeling the Leucines and Valines, and 100 mg/L of methionine [^13^C]. These precursors were added 1hr before induction with IPTG at an OD_600nm_ of 0.6.

### NMR data collection

In addition to the backbone assignment we previously published,^17^ aliphatic and methyl sidechains were assigned using 3D HCCH-TOCSY/CCH-TOCSY, 3D H(CC-CO)NH-TOCSY, and (H)C(C-CO)NH-TOCSY experiments. 3D ^13^C/ ^15^N-edited NOESY^45^ were collected to assign interstrand NOE contacts in both enigma and ground states. A 3D CCH HMQC-NOESY-HMQC was collected with 200ms mixing time on the ILVM-labeled sample to obtain methyl NOE assignment. Methyl NOE contacts were assigned using the methyl side chain assignments and the Alphafold2 structures as a guide.

A modified version of TROSY-based ^15^N constant-time CPMG relaxation dispersion^46^ experiment with relaxation compensation^47^ was performed at 25°C and 35°C at 1.2 mM KaiB^RS^ concentration on a Varian VNMRS DD2 600-MHz system equipped with a triple resonance cold probe. NMR samples contained 1.2 mM KaiB^RS^, 100 mM MOPS, 50 mM NaCl, 2 mM TCEP, and pH 6.5, and 5% D_2_O. Experiments were performed as a pseudo-3D containing 15 2D data sets with a constant-time relaxation period of 60 ms, n_cyc_ values ranging from 0 to 60, 32 scans per FID, and a repetitive delay of 2.5 sec. Repeat experiments were performed with n_cyc_ values of 0 (reference spectrum), 30, and 60.

^15^N-CEST^18^ experiments were recorded on a 800 MHz spectrometer at 4 °C and 20 °C for two different field ^15^N B_1_ field strengths (10 Hz or 20 Hz). B_1_ field calibration was performed as described by ref. ^48^. Data was collected as a pseudo-3D with a relaxation delay, T_relax_, of 500 ms (10 Hz ^15^N B_1_ field) or 400 ms (20 Hz ^15^N B_1_ field), ^15^N offsets ranging between 104 and 135 ppm in increments of 25 Hz, and one reference experiment. Each spectrum was collected with 4 scans per FID and repetitive delay of 1.5 seconds.

Methyl (^13^CH3) ^13^C-CEST^35^ experiment was recorded at 20°C on a 800 MHz spectrometer as a pseudo-3D using a B_1_ radiofrequency field strength of 20 Hz and a relaxation delay, T_relax_, of 400ms. A total of 97 spectra were recorded with the B1 field ranging from 8-25 ppm in steps of 40 Hz. The B_1_ field was calibrated according to the procedure described by ref. ^48^. Each spectrum was collected with 8 scans per FID and a repetitive delay of 2 seconds. The experiment included a reference spectrum recorded with T_relax_ of 0 data set was collected. Methyl (^13^CH3) TROSY-based ^1^H SQ (single quantum) CPMG^36^ experiment (**Fig. S11**) was performed as a pseudo3D containing 22 2D data sets at 20 °C using a constant-time CPMG relaxation period of 40 ms, n_cyc_ values ranging from 1 to 30 Hz. A reference spectrum was collected without a CPMG period (n_cyc_ = 0). Each spectrum was collected with 8 scans per FID and repetitive delay of 2 seconds. Methyl dynamics experiments were performed on 2mM KaiB^RS^ sample in 100 mM MOPS, 50 mM NaCl, 2 mM TCEP, and pH 6.5, and 99% D_2_O.

For real-time NMR, ^15^N CPMG, and ^15^N CEST experiments, peak volumes were integrated in the pseudo-3D dimension using PINT^49^ with a radius of 0.4 ppm and 0.04 ppm in the ^15^N and ^1^H dimensions respectively, and a radius of 0.1 and 0.03 ppm in the ^13^C and ^1^H respectively for the ^13^CH3 CEST and CPMG experiments.

### NMR structural analysis

Secondary structure and order parameters were predicted using TALOS-N^23^. Proline cis/trans probabilities were predicted using PROMEGA^25^. Random coil chemical shifts were predicted using POTENCI^24^. The structure of KaiB-ΔCterm was predicted with CS-Rosetta^43^ using the webserver at https://csrosetta.bmrb.io/ and default settings. The top 10 models from 30,000 decoys are depicted.

### Real-time NMR analysis and fitting

Single-exponential curves were fit to peaks in consecutive HSQC spectra using in-house scripts based on performing a minimization with the lmfit package (https://lmfit.github.io/). Rate constants were fit using a global fit to all assigned Ground or FS peaks that had an amplitude greater 4 x 10^5^ (45 Ground state, 37 FS state peaks). Uncertainty was estimated by bootstrapping where in each iteration, each data point was resampled from the spectrum noise uncertainty, and a new set of residues was resampled with replacement.

### CEST analysis and fitting

We first identified visually from CEST data which residues had 2 or 3 states present at both 4°C and 20°C, taking into consideration assignments of the Ground and PD states. For each of those residues, we deductively reasoned based on known PD assignments which peaks corresponded to which states. We fit 2-state and 3-state models in ChemEx^18^. For 3-state fits, we constrained R_2_, i.e. the transverse relaxation rate, of each residue of the Enigma state to match its R_2_ of the Ground state. Exchange with the PD state was most observable at 4°C, and 11 residues were used to fit a 3-state model at 4°C. At 20°C, we could fit 5 residues to a 3-state model. We found that the Enigma-PD interconversion was underdetermined due to the low populations of both, and found that other models had comparable reduced chi^2^ values. For instance, allowing interconversion between the Enigma and PD state, i.e. all 3 states fully connected, resulted in comparable fits with Enigma ↔ PD exchange on the order of 1e-6 /s. Our data therefore does not rule out the possibility of interconversion between the Enigma and PD state – because both are minorly populated, this transition would be unobservable in CEST, meaning the fit is underdetermined in respect to this one transition. Alternate plausible models are depicted in **Fig. S5**. Allowing all states to interconvert, and constraining *k_Ground->PD_ = k_Enigma->PD_* resulted in comparable reduced Chi^2^ models, but removing the Ground-PD interconversion results in notably worse models by reduced Chi^2^. Taken together, this suggests that there could be even up to equal flux to the PD state through the enigma state as directly from the Ground state, though the path cannot be exclusively through the enigma state.

In the ^13^CH3 CEST data (**Fig. S12**), most of the profiles have 2 dips indicating exchange between 2 states and a few have 3 states indicating the presence of a third state, similarly to the ^15^N-CEST. Residues with 2 states in the ^13^CH3 CEST data were selected for analysis. We fit these residues to a 2-state exchange model using ChemEx, with *k_e”_* = 173 ± 8 s^-1^ and *p_b_* = 7 ± 1 %, which is in agreement with the rate and population obtained from ^15^N-CEST. There were about 5 profiles that show 3 states, but we focused on the 2 state exchange for the methyls in order to assign the enigma state. Similarly to the ^15^N-CEST, the major state dip matches the assignment of the ground state which is the most populated at 20 °C. The majority of profiles have small Δω_c_ of < 0.7 ppm (**Fig. S12**).

#### Methyl CPMG analysis

R_ex_ was estimated for the Ground state methyl peaks as *R_2,eff_*(*ν*=25 Hz) – *R_2,eff_*(*ν*=∞), where *R_2,eff_*(*ν*=∞) using the ^13^CH3 TROSY-based ^1^H SQ data. Inspection of the CPMG profiles show that two groups of methyl resonances are undergoing exchange with different rates. The majority of the residues with large R_ex_ > 5 s^-1^ are clustered around the register shifted strand. All methyl CPMG data are shown in **Fig. S13**. Amongst these residues, L82, V81, L85, I54, I57, and I72 have larger Δω_C_ with L82 showing the highest Δω_C_ of 4.1 ppm. The observed Δω correlates with the difference in the enigma and ground state structures. For example, L82 is in close contact with L85 in the ground state and facing the hydrophobic patch, but in the enigma state, L82 points outward and a slight rearrangement of the loop brings I54 closer to it. Meanwhile in the ground state, I52 is too far to observe an exchange cross-peak with L82 but instead we observe an exchange cross-peak between L82 and I57 which supports the structural difference between the 2 states (cf. **Fig. S10**). Another example is the exchange cross-peak we see between L85 and L60. In the enigma state, the register shift changes the position of L85 relative to L60, thereby slightly changing the environment of both methyls (cf. **Fig. S10**). Using a combination of Δω_C_ from methyl-CEST, and CCH-methyl NOESY experiments, we were able to assign these to the enigma state.

### CPMG analysis and fitting

Peak volumes were integrated in the pseudo-3D dimension using PINT^49^ with a radius of 0.4 ppm in the ^15^N dimension and a radius of 0.04 ppm in the ^1^H dimension. Uncertainty was estimated in the noise level in the spectrum (using the –noiseUncertainty flag in PINT^49^). Two duplicate data points were collected. The larger uncertainty of either the noise-based or duplicate-based uncertainty was reported.

CPMG data was collected at two temperatures, 25°C and 35°C, to assess if each peak with noticeable elevated *R_2,eff_* was experiencing exchange in the fast or slow CPMG regime, as data collected at one field strength or temperature alone is insufficient to determine this^28,29^. We refer the reader to ref. ^28^ for a more complete theoretical treatment. To briefly understand the theoretical basis for this, consider a two-state system of exchange between states A and B, with populations given by *p_A_* and *p_B_*, and forward/reverse rates given by *k_AB_* and *k_BA_*. Let 𝑅_2,𝐴’_ be the observed transverse relaxation rate of state A. If exchange is occurring in the fast limit, then 𝑅_2,𝐴’_ may be approximated as

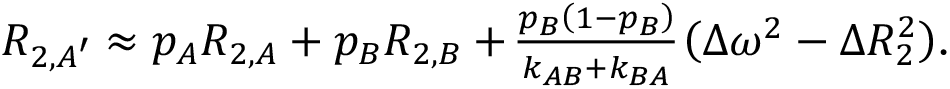

An increase in temperature would cause an increase in *k_AB_+k_BA_*, which would decrease *R_ex._* Note that 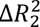 can be assumed to be small in comparison to Δ𝜔^2^.

If exchange is occurring in the slow limit, then *R_ex_* isis essentially the rate of leaving the observed state, i.e.

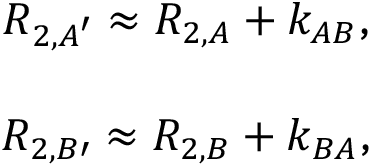

Where 𝑅_2,𝐴_ and 𝑅_2,𝐵_ are the intrinsic transverse relaxation rates of states *A* and *B.* An increase in temperature means that *Rex* will increase. Note this means that the modified Carver-Richards equation cannot be used to infer p_B_ and k_ex_ values.^28^

For each Ground state peak, *R_ex_* was estimated as *R_2,eff_*(*ν*=50 Hz) – *R_2,eff_*(*ν*=∞), where *R_2,eff_*(*ν*=∞) is estimated as the mean of the last two collected data points. *R_ex_* at 25°C and 35°C were compared with a nonparametric significance test to determine if *R_ex_* at 35°C was higher or lower than that at 25°C to p<0.05. A Bonferroni multiple hypothesis correlation was applied. The resulting *R_ex_* values and significance tests are in **Supplemental Data 2**. Based on this analysis, the majority of the residues were in the slow regime.

### AF2 structure predictions

KaiB models from AF-Cluster were taken from the data repository at https://github.com/HWaymentSteele/AFCluster/. Single-sequence AF2 sampling was performed in ColabFold^50^ using ‘msa_mode=single_sequence’, dropout=True, num_seeds=16, and all other options set to default.

### Chemical shift prediction and AF2 model analysis. SPARTA+ was run using the command

‘sparta+ -i <input.pdb>’ on NMRbox. SHIFTX2 was run using the webserver at http://www.shiftx2.ca/ with pH=6.5 and temperature=293 K and default settings otherwise. UCBshift was run on NMRbox using the command ‘CSpred -b single_seq_files -o ss_outputs -x -pH 6.5’ where <LIST_OF_PDBS> is a list of the pdb models for which to generate predictions. pLDDT was calculated as the mean pLDDT over C_a_ atoms. Spearman correlations between actual and predicted chemical shifts were calculated using SciPy^51^.

### Free energy and barrier height estimation from real-time NMR and CEST populations

To combine population estimates of the FS state from real-time NMR with population estimates from CEST, in which the FS state is unobservable, we re-scaled the populations estimated with CEST (the Enigma, Ground, and FS state) to the overall population of the Ground state estimated from real-time NMR.

We calculated barrier height of the transition from state *A* to *B,* Δ𝐺^‡^ using the Eyring equation,

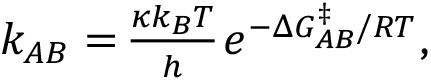

rewritten to solve for Δ𝐺^‡^:

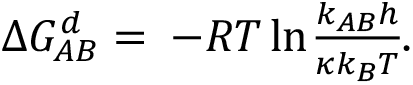

Here, *k_AB_* is the measured interconversion rate from *A to B,* 𝜅, the transmission coefficient, is assumed to be 1, *k_B_* is Boltzmann’s constant, T is temperature in Kelvin, *h* is Planck’s constant.

We estimated an absolute offset between Δ𝐺^𝐺𝑟𝑜𝑢𝑛𝑑^(4°𝐶) and Δ𝐺^𝐺𝑟𝑜𝑢𝑛𝑑^(20°𝐶) by assuming the absolute barrier height of a reaction cannot decrease with temperature. We therefore added a constant offset of 2 kcal/mol to Δ𝐺 values originally estimated at 4°C to account for this. All the above calculations are included in **Supplemental Data 3**.

### FoldX prediction

We predicted energies for both the Ground and Enigma structures using FoldX^31^, and inspected the energy terms related to hydrogen bonding, electrostatics, penalty from burying polar residues, and stabilization from burying hydrophobic residues. Predicted hydrogen-bonding and electrostatic energies were very similar between both structures. The largest stabilizations of the Ground state over the Enigma state came from van der Waals energy and energy of burying hydrophobic charges.

We noticed two possible driving forces for the Enigma state. The first is from four charged residues: Asn84, Lys86, Asp88, and Glu90 in the C-terminus. These residues are solvent-facing in the FS state, but are buried in the core of the Ground state. The register shift of the Enigma structure allows these four charged residues to be more solvent-exposed. FoldX predicts to gain roughly 1 kcal/mol of stabilization each in the Enigma state when compared to the Ground state.

Another possible driving force comes from a hydrophobic patch present on the Ground state structure, formed by residues Ile54, Val55, and Leu82. In KaiB from *T. elongatus*, this hydrophobic patch forms the dimerization interface of the Ground state. The Enigma state abolishes this hydrophobic patch: Leu82 switches from being displayed on the surface to buried in the hydrophobic core. Val81 is instead on the surface of the protein, but the register shift causes it to shift such that Ile54 and Val55 can cover it. The magnitude of this is expected to be smaller than the first possible Enigma driving force, with only 2 kcal/mol energy difference across all energy terms for these 4 residues.

## Data availability

Chemical shifts for KaiB^RS^ are being made available at BMRB accession number 52018. Raw data of real-time NMR experiments, relaxation dispersion experiments, structure models, and scripts for single-exponential fits are available at https://zenodo.org/doi/10.5281/zenodo.11448116.

## Code availability

Code for performing the CPMG error analysis described here are available at https://github.com/HWaymentSteele/NMR_scripts.

## Notes

https://zenodo.org/doi/10.5281/zenodo.11448116

### Summary of Updates

Updated version containing further experimental results.

