## Supporting information for "The conformational landscape of fold-switcher KaiB is tuned to the circadian rhythm timescale"

**This PDF file includes:**

Figures S1 to S15
Tables S1
Captions for supplemental data 1-3
SI References

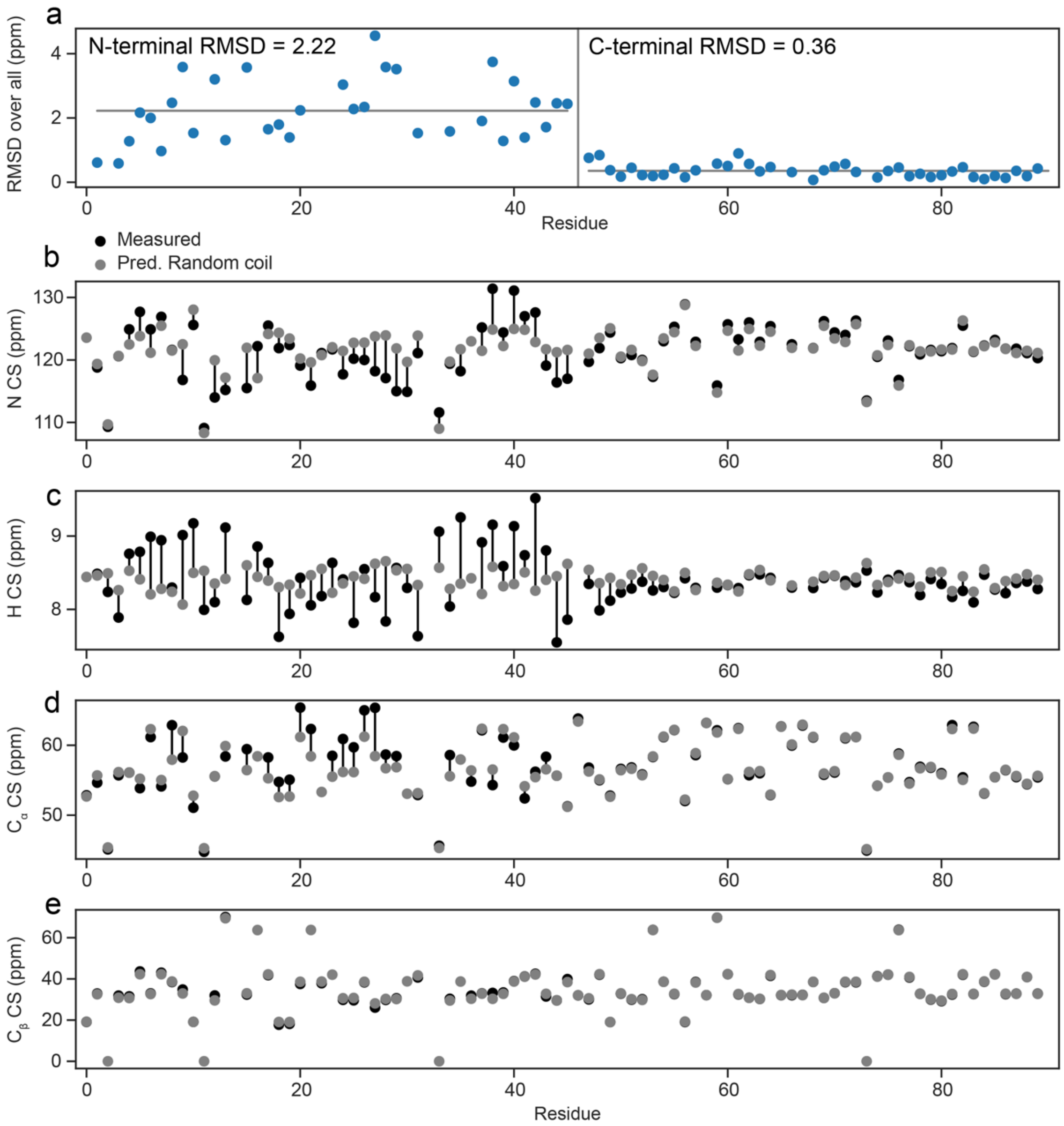


**Fig. S1. Major state at 4˚C is a partially disordered state.** Chemical shifts of C-terminus (Leu46 onwards) for major state at 4˚C correlate well with random coil chemical shifts predicted by POTENCI[1]. (a) RMSD per residue of N, H, Cα, Cβ chemical shifts relative to random coil shifts. (b-e) Measured (black) vs. predicted (grey) for random coil chemical shifts for N, H, Cα, Cβ atoms.

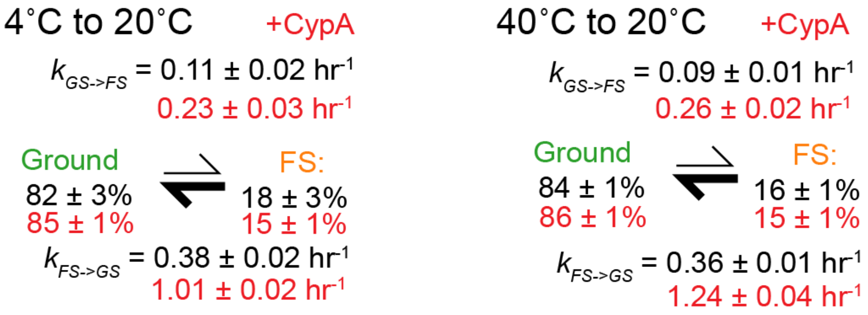


**Fig. S2. Kinetics for Ground to FS interconversion and rate acceleration by CypA.** Rate constants and populations fit from KaiB incubated at 4˚C (left) or 40 ˚C (right) and monitored at 20˚C (compare to Fig. 1b).

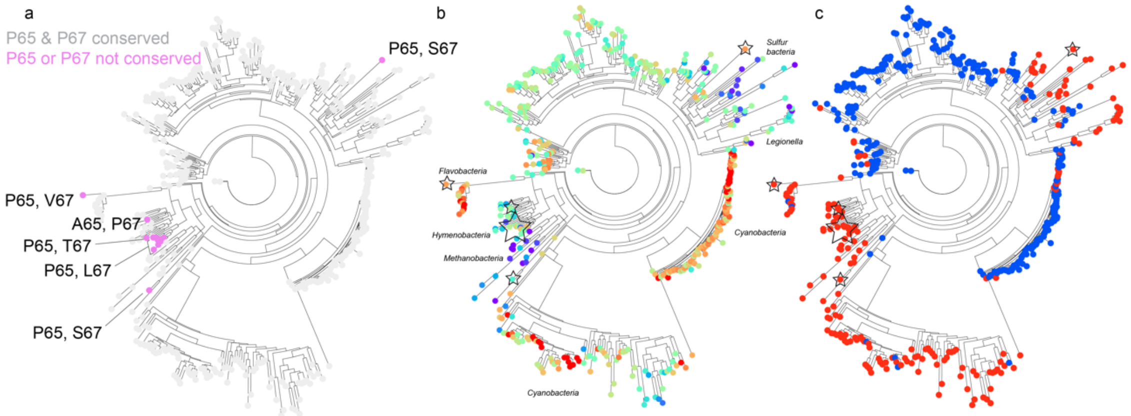


**Figure S3. Conservation of P65 and P67 across KaiB phylogenetic tree** in ref. [2]. a) Sequences in grey have P65 and P67 conserved. Sequences in magenta have only one of these two prolines, mutations noted on figure. b) Tree colored by pLDDT from closest-10 AF2 predictions and c) Predicted FS (red) or Ground state (blue) from ref. [2].

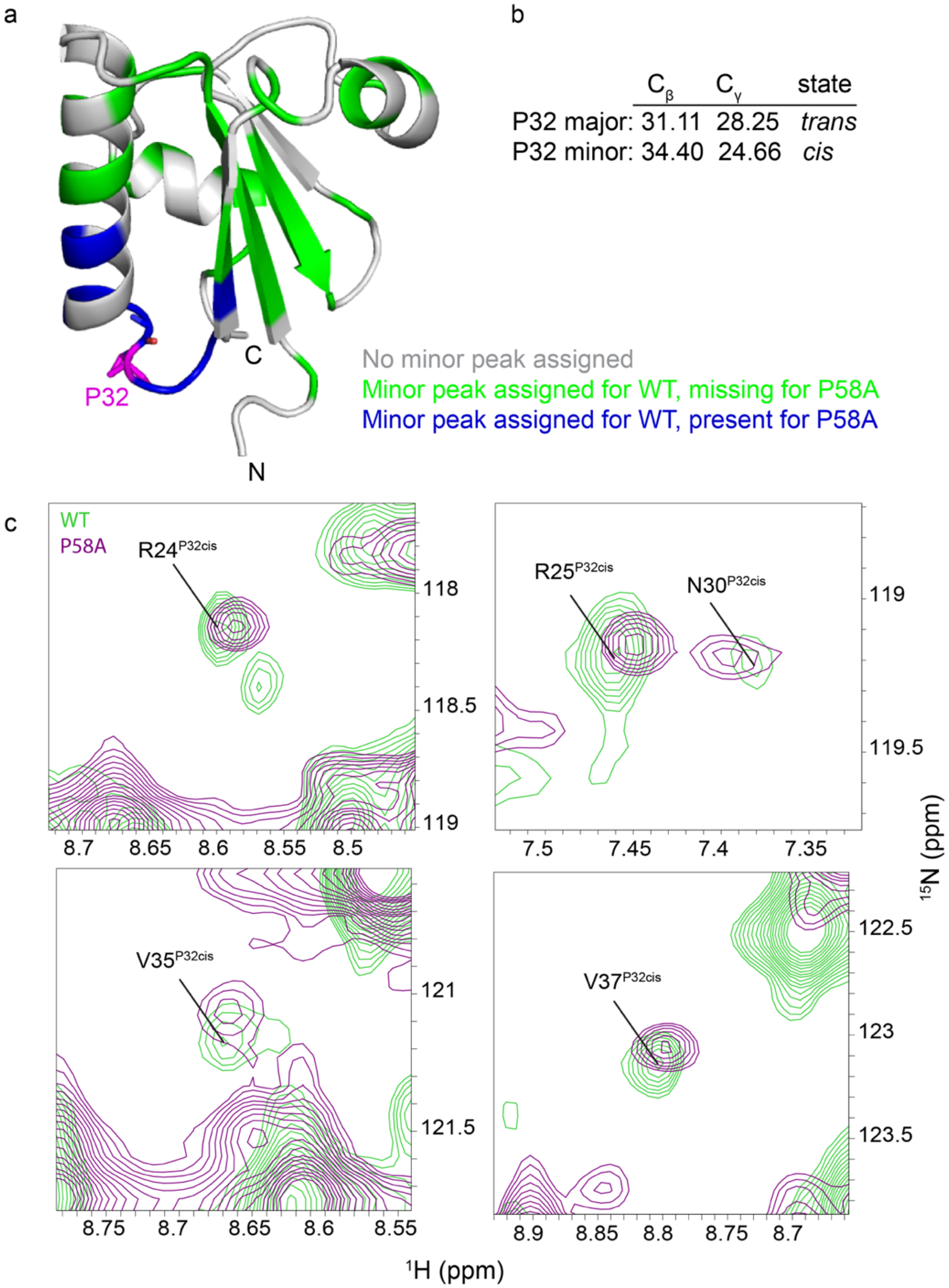


**Fig. S4. Some minor peaks correspond to a minor *cis* state of P32.** a) Fold-switched structure of KaiB from AF-Cluster, colored by grey: no minor peak could be assigned, green: a minor peak was assigned in KaiB WT but not in KaiB P58A, blue: a minor peak was assigned in both KaiB WT and KaiB P58A. b) The $C_\beta$ and $C_\gamma$ shifts of the major and minor peak of P32 correspond to a trans and cis state. c) representative minor peaks present for both KaiB WT (green) and KaiB P58A (purple).

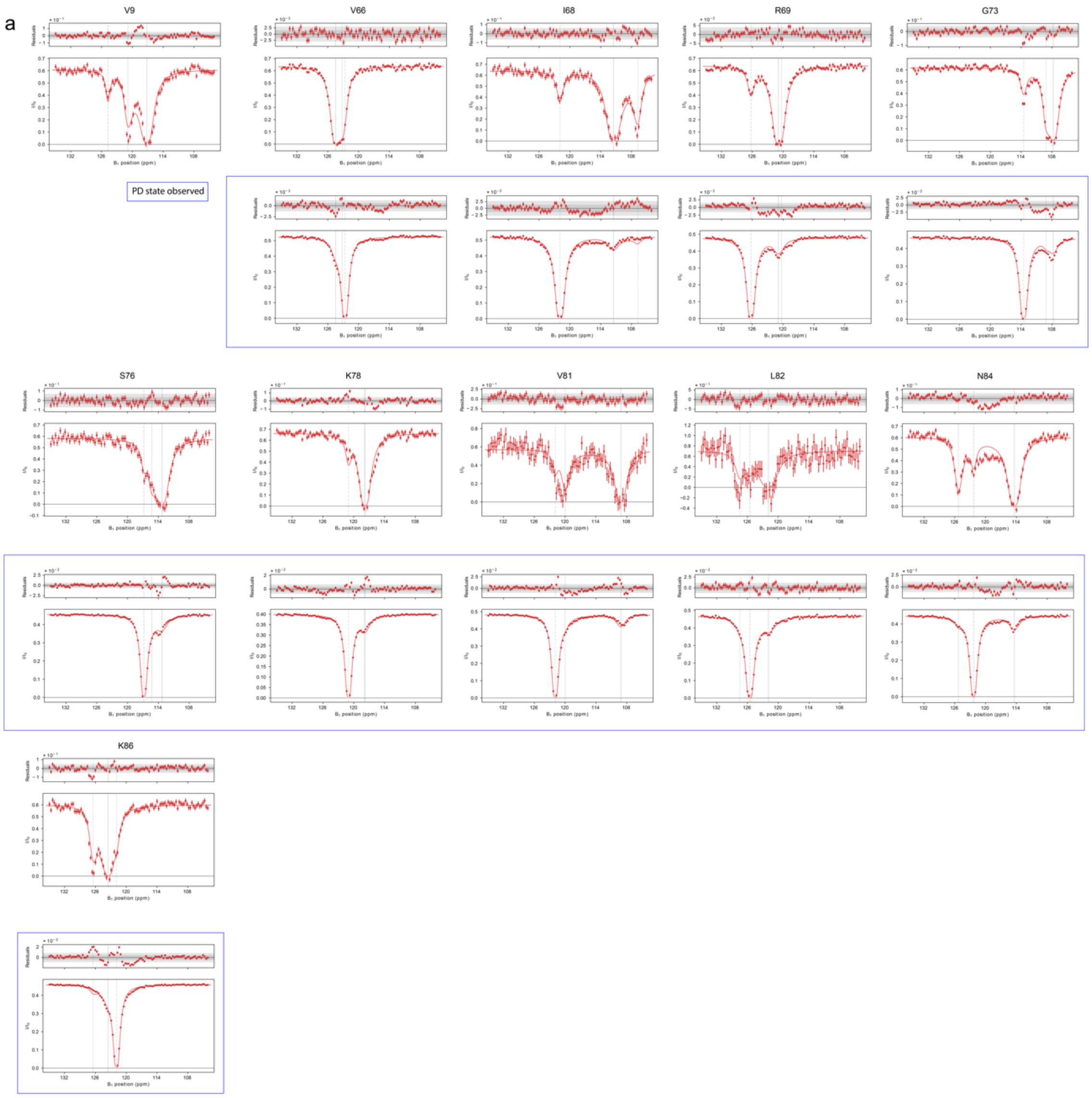
a
V9
V66
I68
R69
G73
PD state observed
S76
K78
V81
L82
N84
K86
Residuals
B₁ position (ppm)

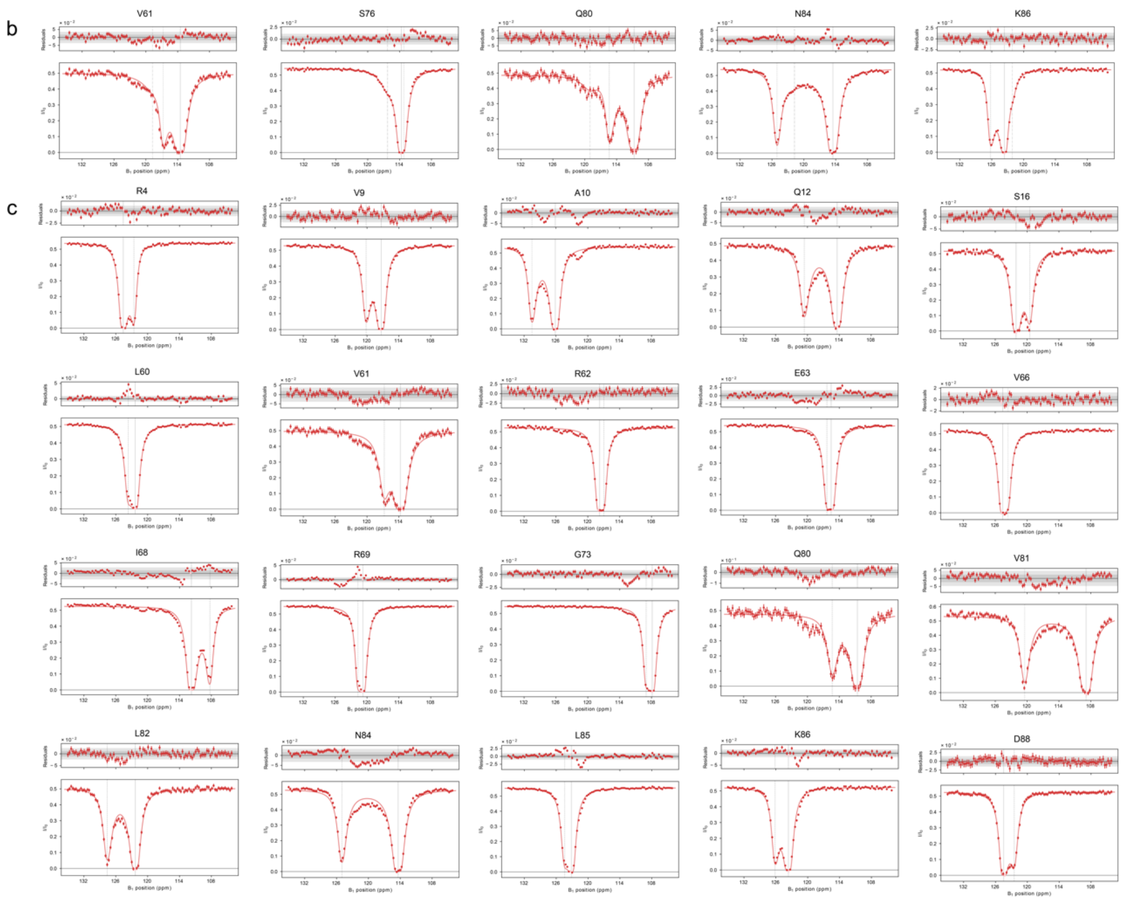


**Fig. S5 (multi-page figure).** **$^{15}$N CEST data overlaid with global fits** for (a) 3-state fit at 4˚C (PD state observed is boxed in blue), (b) 3-state fit at 20˚C, (c) 2-state fit to Ground and Enigma state at 20˚C.

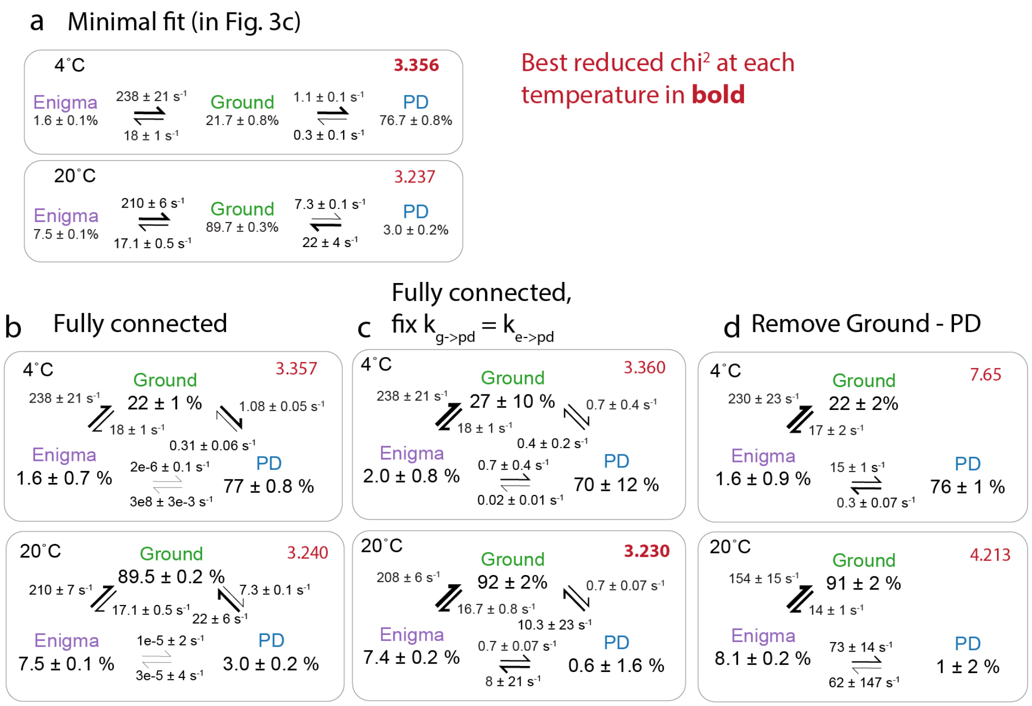


**Fig. S6. Evaluating alternate models for fitting CEST data.** (a) Minimal models representing data, reproduced from Fig. 3c. (b) Allowing all states to interconvert. (c) Constraining $k_{Ground\text{->}PD} = k_{Enigma\text{->}PD}$. (d) Removing the Ground-PD interconversion results in notably worse models by reduced $Chi^2$. Taken together, this suggests that there could be even up to equal flux to the PD state through the enigma state as directly from the Ground state. The enigma to PD interconversion is underdetermined because of the too low populations. Physically, it would make most sense if both GS and enigma can interconvert with the PD state given their similar structures. These alternate fits rule out the path going exclusively through the Enigma state.

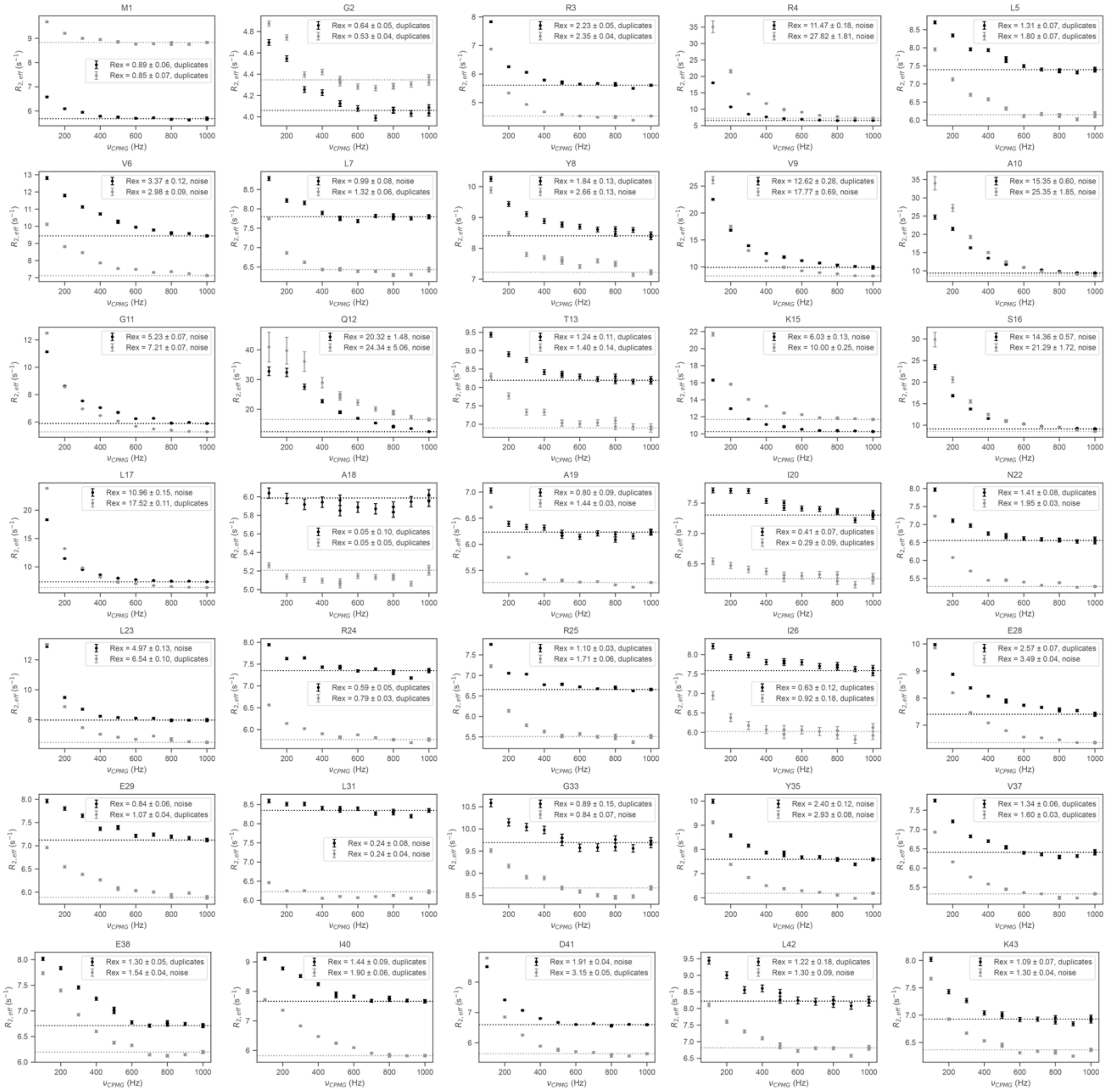

**Fig. S7. (multi-page figure, 1 of 2). $^{15}$N CPMG data for all assigned Ground state peaks at 25˚C and 35˚C.**

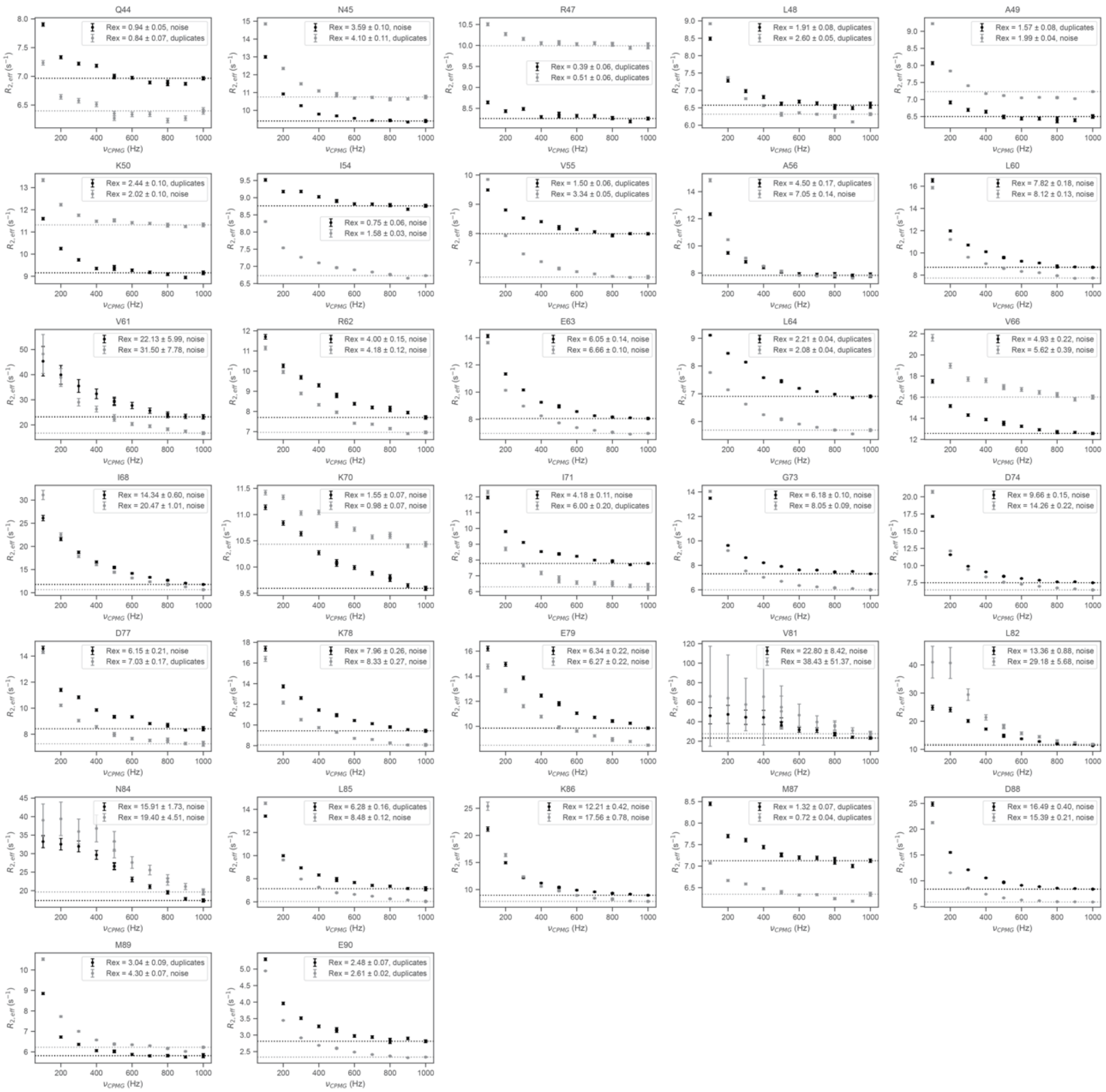


**Fig. S7. (multi-page figure, 2 of 2).** $^{15}$N CPMG data for all assigned Ground state peaks at 25˚C and 35˚C.

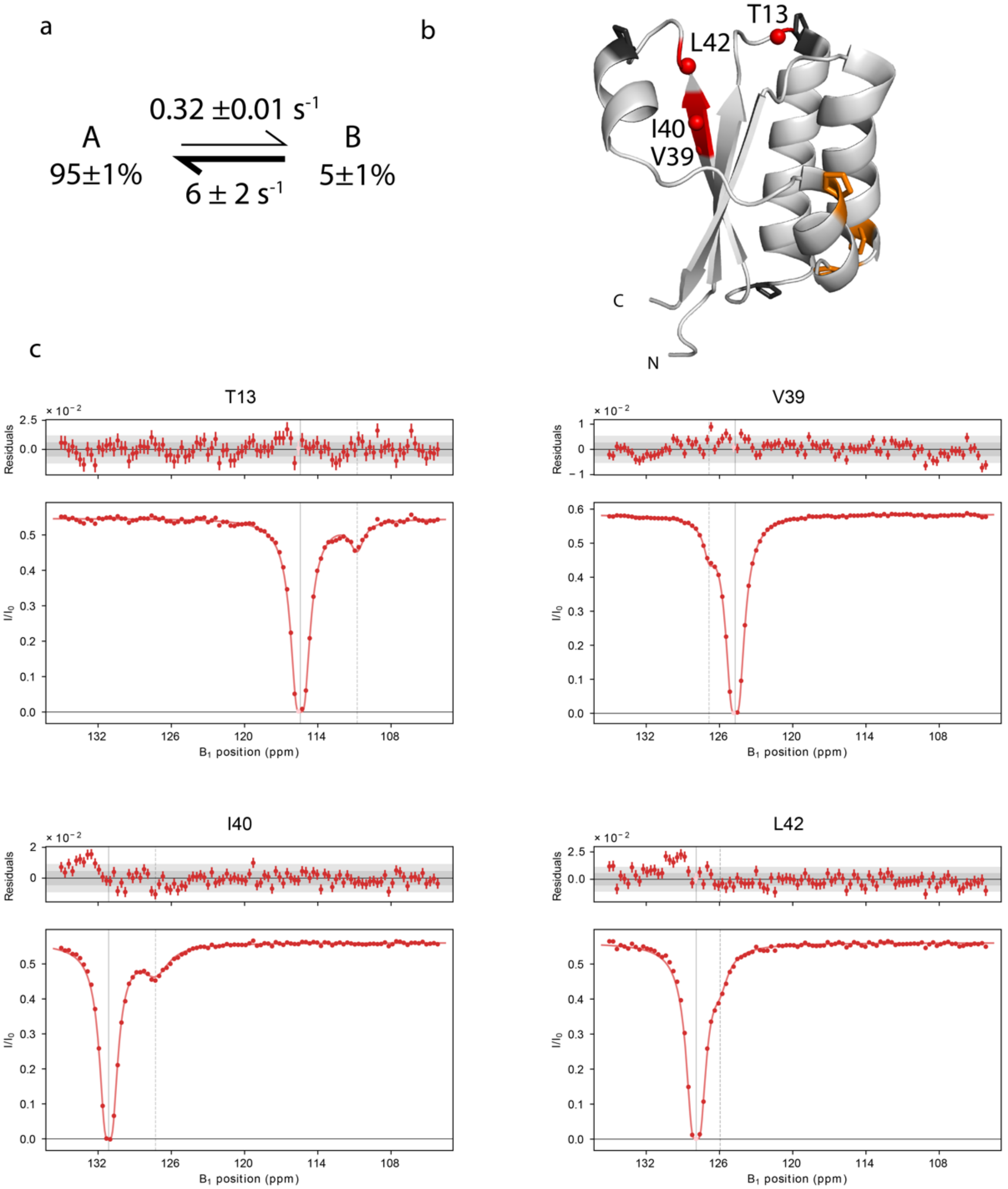


**Fig. S8. Different minor slow process fit in CEST data at 20˚C, likely due to cis/trans prolyl isomerization of Pro14**. a) Populations and kinetics from 2-state fit. b) Location of residues fit to process visualized in red. Prolines P14, P32, P48 shown in black. Prolines P58, P65, P67, which are *trans* in the Ground and *cis* in the FS state, are shown in orange. c) CEST data from field strength = 20 Hz overlaid with fits from ChemEx.

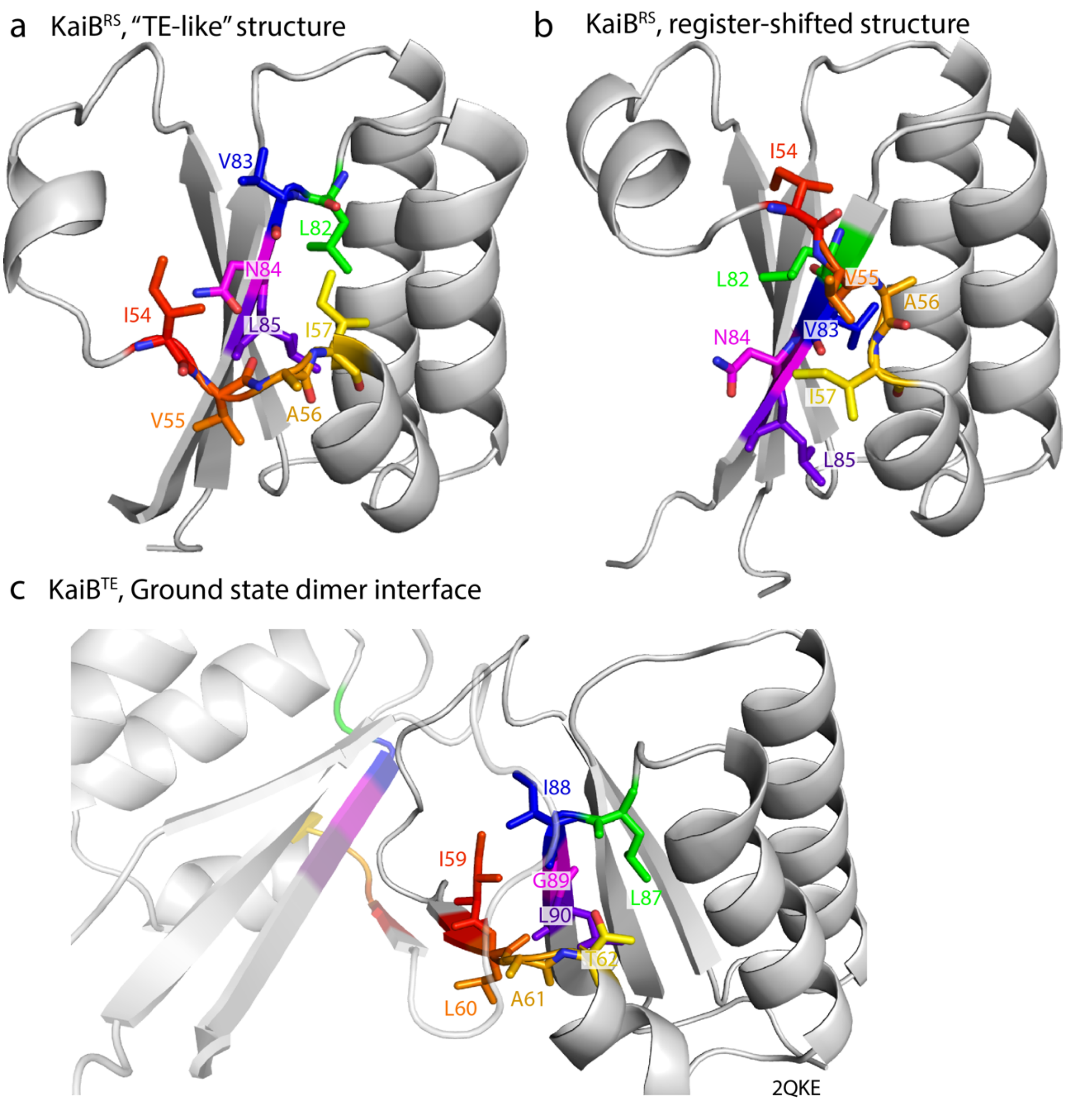


**Figure S9**. Difference in hydrophobic packing of core between (a) TE-like and (b) register-shifted, proposed Enigma structure. (c) The hydrogen-bonding and hydrophobic core of the TE-like structure is homologous to the crystal structure of KaiB from *Thermosynechococcus elongatus* (PDB: 2QKE[3]). I59, L60, A61 are involved in the dimerization interface in the tetrameric structure in *T. elongatus*, and this same geometry is predicted for homologous residues in the TE-like structure of *R. sphaeroides* (compare to (a)).

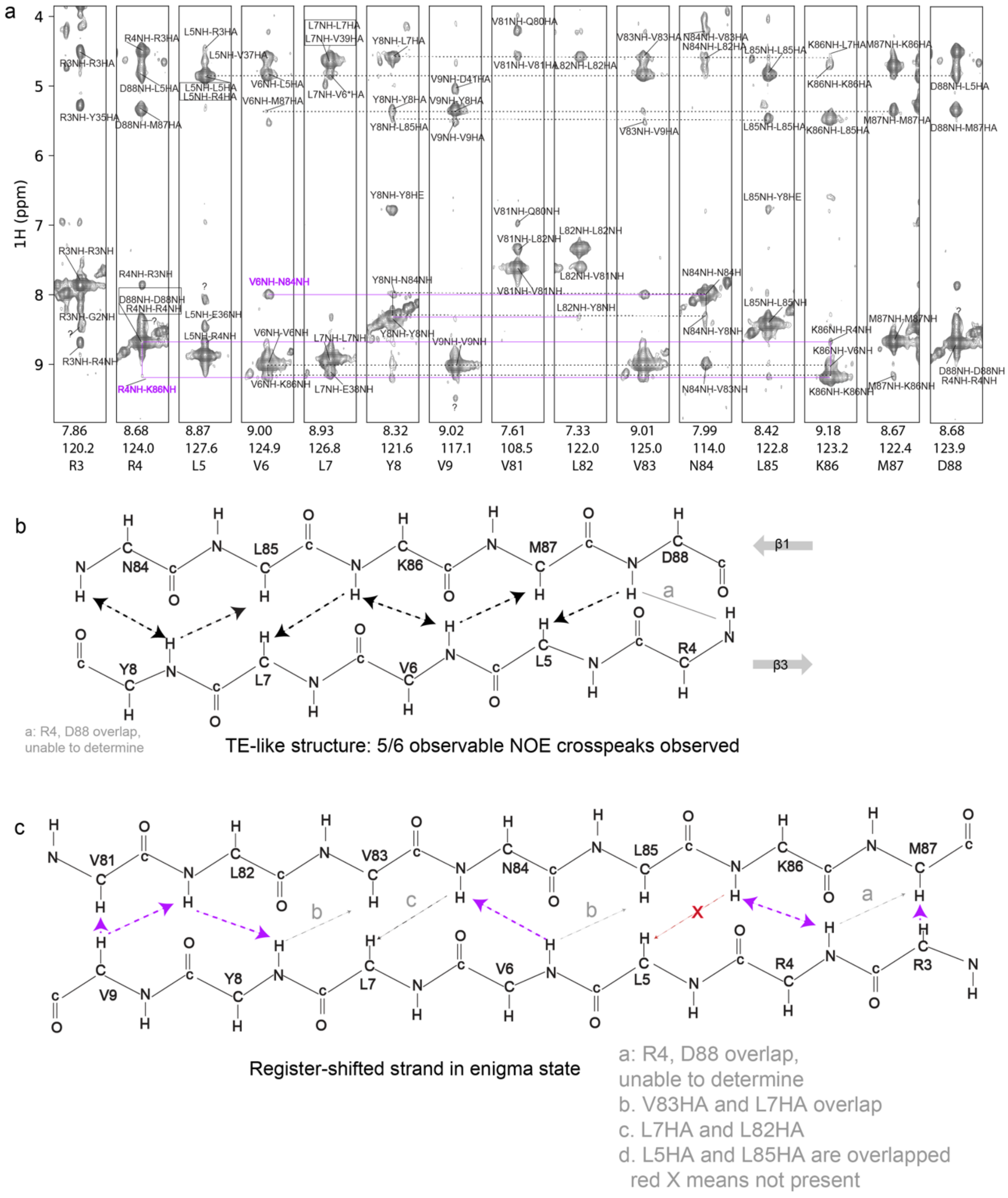


**Fig. S10 (2-page figure). NOESY data supporting the TE-like structure as the major ground state, and minor enigma state as register shifted conformation.** a) $^{15}$N-edited NOESY supporting inter-strand NOE cross-peaks for both TE-like and register-shifted structure. TE-like NOE cross-peaks are annotated in black and Register-shifted cross-peaks are annotated in purple. b) Scheme of identified cross-peaks supporting TE-like structure. c) Scheme of identified cross-peaks supporting register-shifted structure. In (b) and (c), gray arrows indicate cross peaks where due to overlap, inter-strand NOE cross-peaks are ambiguous while red arrow indicates missing NOE cross-peak.

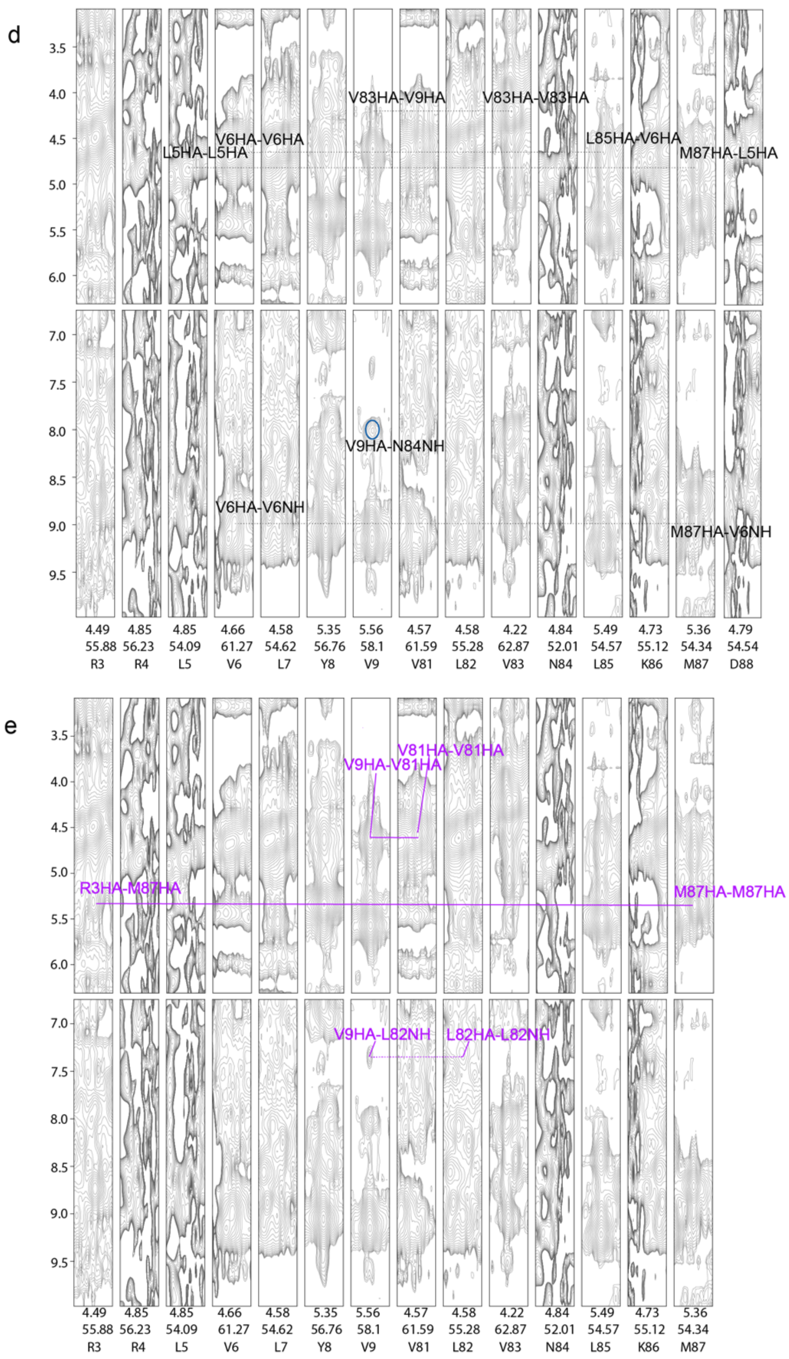


**Fig. S10 (2-page figure). NOESY data supporting the TE-like structure as the major ground state, and minor enigma state as register shifted conformation.** d) $^{13}$C-edited NOESY annotated with cross-peaks supporting TE-like structure in black. e) $^{13}$C-edited NOESY annotated with cross-peaks supporting register-shifted structure in purple.

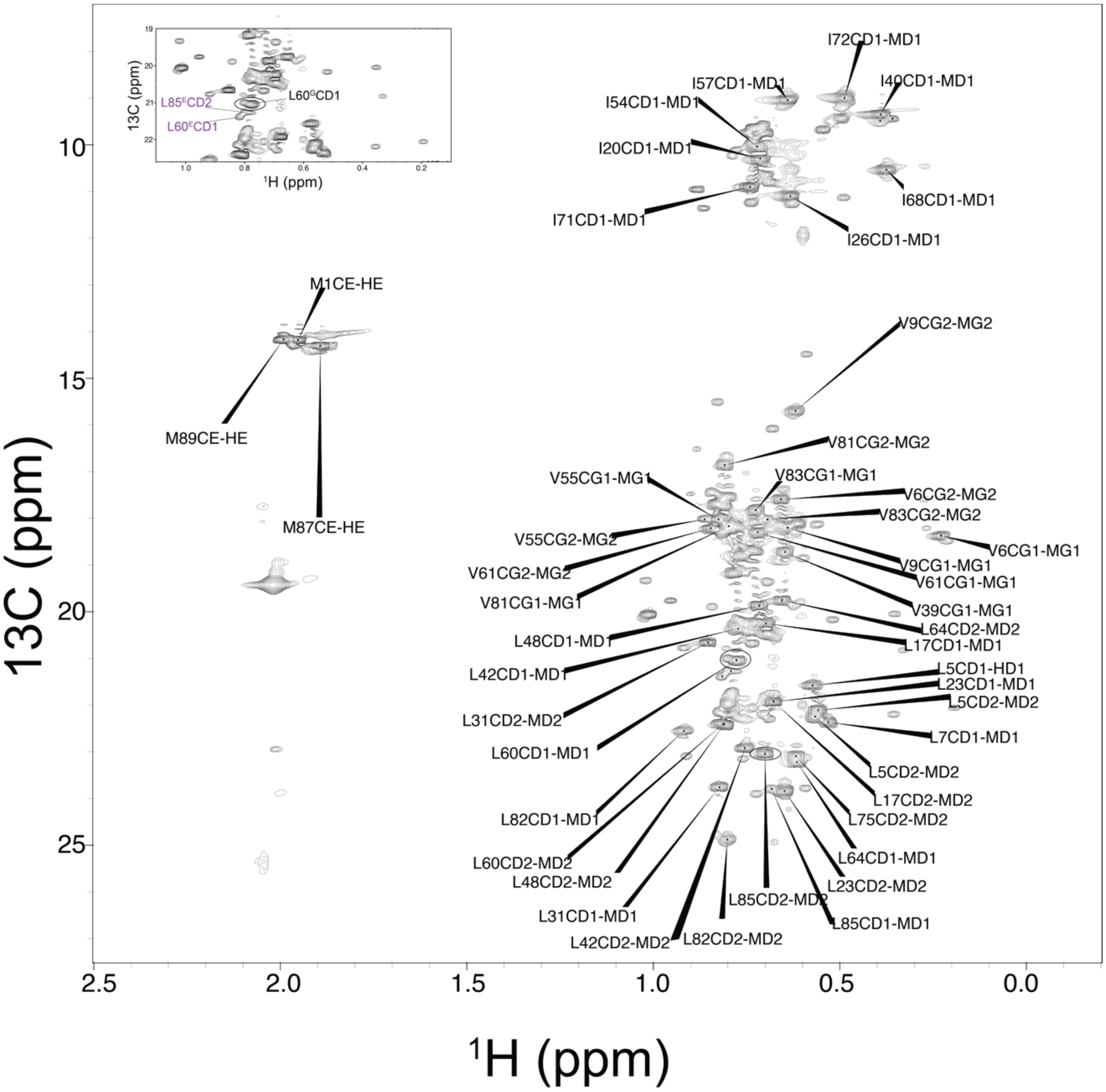


**Fig. S11**. $^{13}$C-$^{1}$H HMQC 2D collected at 20˚C on $^{13}$C ILVM methyl-labelled KaiB. Ground state assignments are shown. Inset shows zoomed in region where L60 and L85 enigma state peaks are indicated in purple labels. Their corresponding ground state peaks are circled.

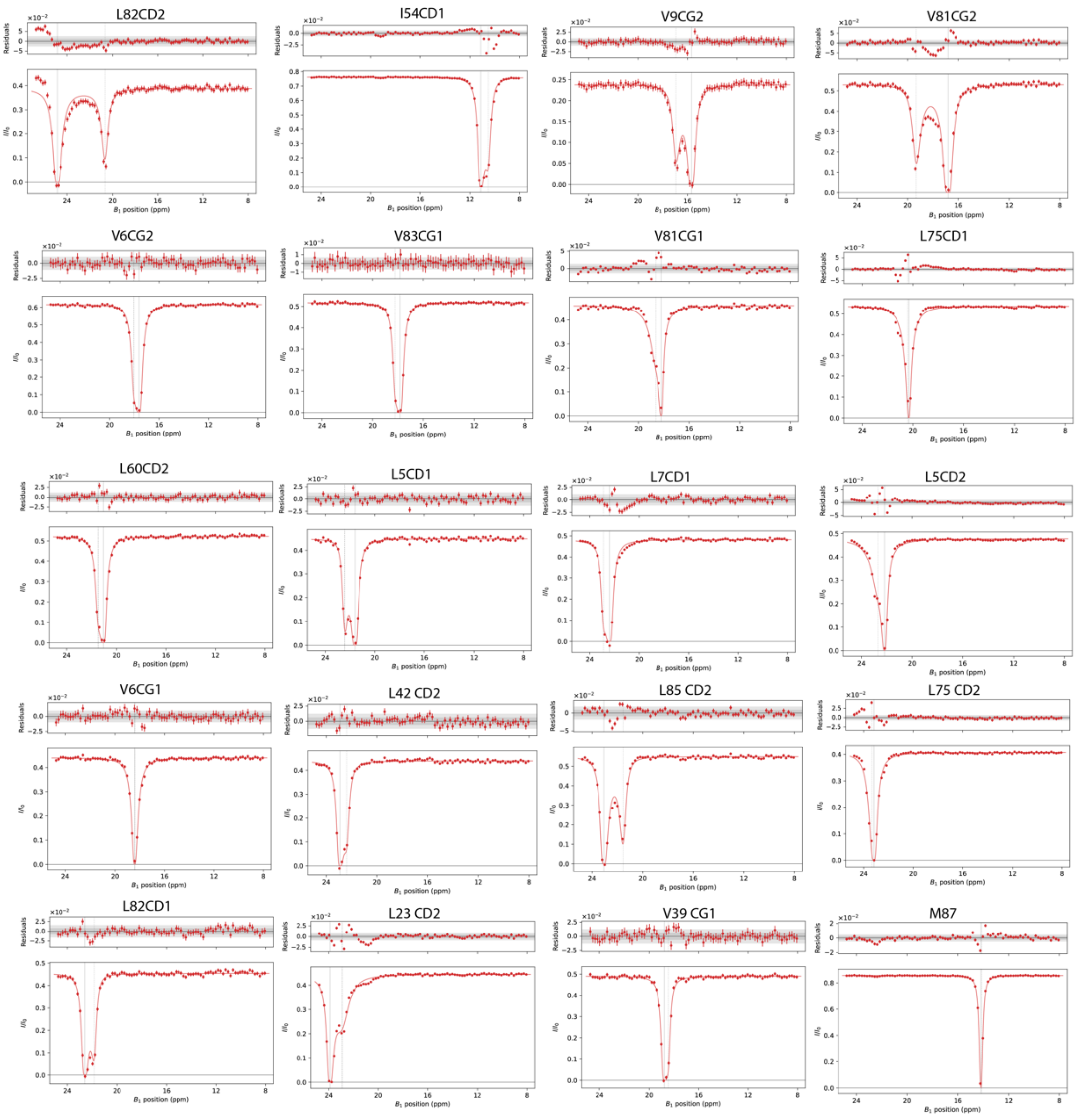


**Fig. S12. $^{13}$C-$CH_3$ CEST data at 20˚C overlaid with global 2-state fit between Ground and Enigma state.** This fit obtained $k_{ex}$ = 173 ± 8 $s^{-1}$ and $p_b$ = 7 ± 1%, in agreement with the rate and population obtained from $^{15}$N-CEST.

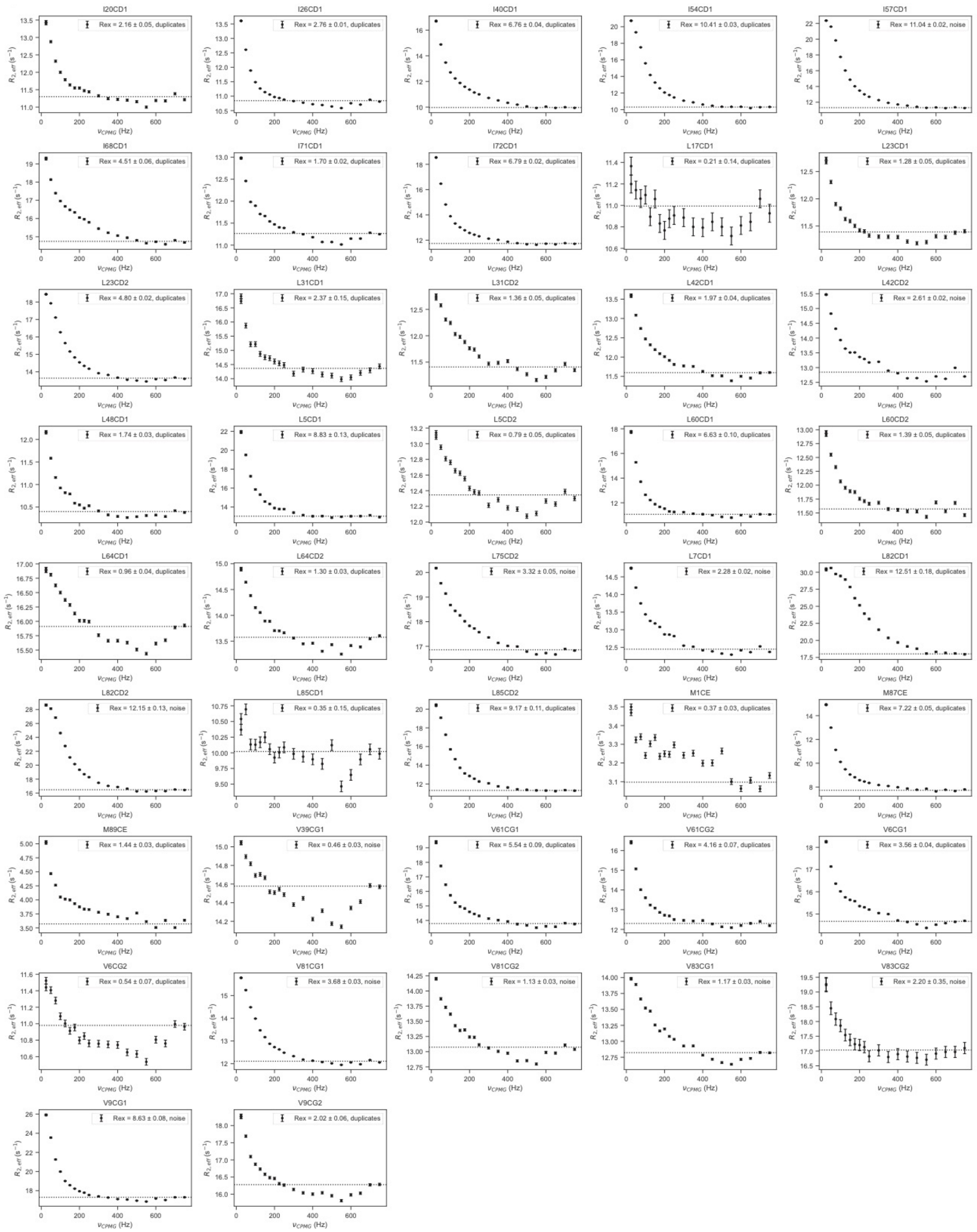


**Fig. S13.** All $^{13}$C-CH3 CPMG data at 20 ˚C.

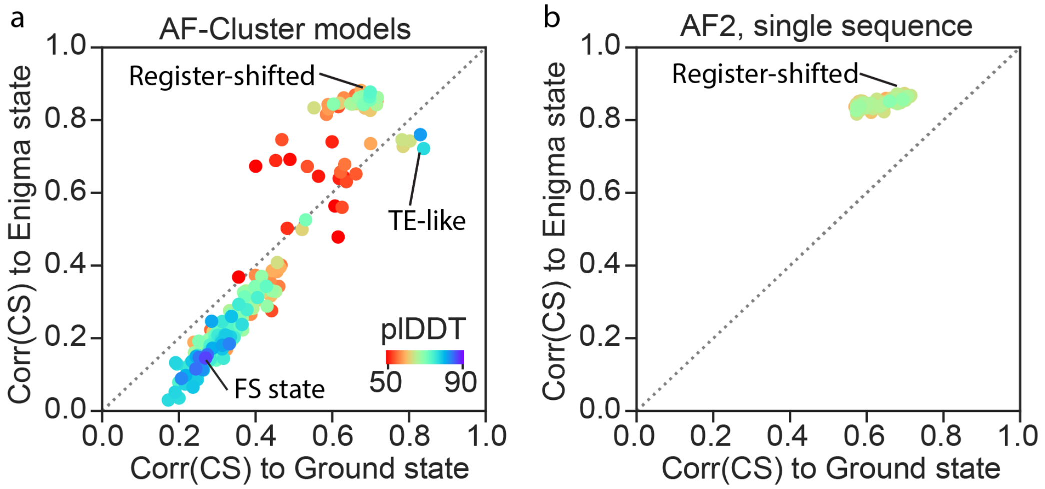


**Figure S14. AF2 in single-sequence mode predicts the Enigma state**. Correlation of $^{15}N$ chemical shifts predicted using UCBshift[4] with Ground and Enigma chemical shifts, plotted for all KaiB structure models generated by (a) AF-Cluster[2], and (b) increased sampling of AF2[5] in single-sequence mode. AF-Cluster models show difference in correlation to either chemical shifts from the Ground or Enigma state, and AF2 single-sequence sampling only samples the register-shifted structure.

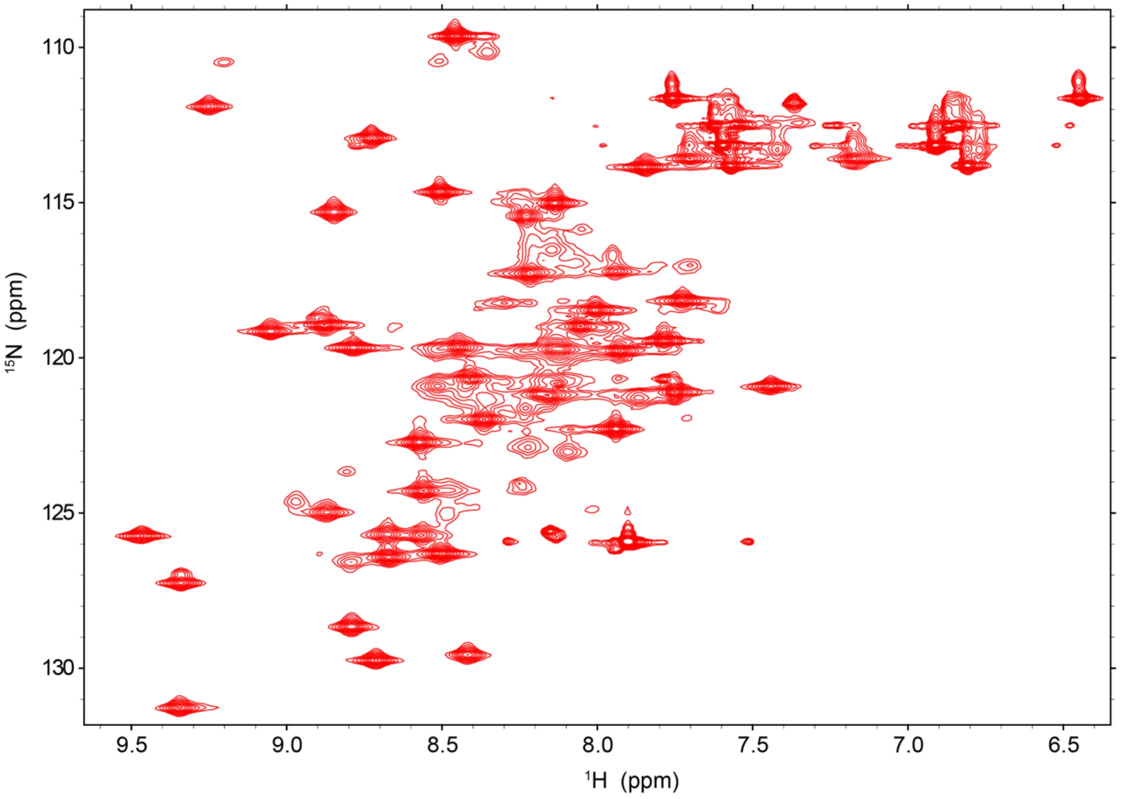


**Figure S15. KaiB-ΔCterm is folded over temperature range from 4 ˚C to 25 ˚C.** $^{1}H$-$^{15}N$ HSQC of KaiB-ΔCterm at 25˚C.

**Table S1.** Proline isomerization states in homologous structures of KaiB Ground and FS state. "—"= proline not conserved.

| State | PDB code | Method | Organism / notes | Reference | T13-P14 | L31-P32 | N45-P46 | I57-P58 | L64-P65 | V66-P67 |
|---|---|---|---|---|---|---|---|---|---|---|
| Ground | 2QKE | X-ray | *T. elongatus* | Pattanayek et al. (2008)[3] | *trans* | *trans* | *trans* | *trans* | *trans* | *trans* |
| Ground | n/a | NMR | *R. sphaeroides* | This work | *trans* | *trans/cis* | *trans* | *trans* | *trans* | *trans* |
| FS | 4KUN | X-ray | *L. pneumophila* | Loza-Correa et al. (2014)[6] | -- | *trans* | -- | *cis* | *cis* | *cis* |
| FS | 5JYT | NMR | *T. elongatus* + mutations | Tseng et al. (2017)[7] | *trans* | -- | *trans* | *cis* | *cis* | *cis* |
| FS | 8FWJ | Cryo-EM | *R. sphaeroides* | Pitsawong et al. (2023)[8] | *trans* | *trans* | *trans* | *cis* | *cis* | *cis* |
| FS | 8UBH | NMR | *T. elongatus vestitus,* KaiB-4 | Wayment-Steele et al. (2024)[2] | -- | *trans* | *trans* | *cis* | *cis* | *cis* |
| FS | n/a | NMR | *R. sphaeroides* | This work | *trans* | *trans/cis* | *trans* | *cis* | *cis* | *cis* |
| PD | n/a | NMR | *R. sphaeroides* | This work | *trans* | *trans/cis* | *trans* | *trans* | *trans* | *trans* |

**Supplemental Data 1. Energetic contributions for TE-like and register-shifted structures estimated with FoldX[9].**

**Supplemental Data 2. Estimated $R_{ex}$ and determined exchange regime from CPMG at 25˚C and 35˚C.**

**Supplemental Data 3. Calculations pertaining to barrier heights and populations of all states for KaiB depicted in Fig. 5.**
